# Elevated isoform richness in males largely reflects transcriptional noise rather than proteomic complexity

**DOI:** 10.64898/2026.08.10.744030

**Authors:** Linley M. Sherin, Bernadette D. Johnson, Alberto Corral-Lopez, Wouter van der Bijl, Judith E. Mank

## Abstract

Alternative splicing (AS) can generate multiple RNA isoforms from a single gene and is thought to contribute to phenotypic divergence, including differences between the sexes. Studies in several organisms have documented sex differences in splicing, however these have been largely reliant on short-read RNA sequencing which requires complex algorithms to assemble full-length transcripts and may underestimate both isoform diversity and sex differences in splicing. We used long-read, single molecule RNA-Seq to build a more complete catalog of sex-biased splicing in *Poecilia reticulata,* a focal species for studies of sexual dimorphism. Pairing long-read sequencing with isoform-level analyses, we identified a sixfold higher proportion of sex-biased splicing genes (37%) compared with short and long-read event-level approaches (6%, 11%). AS was common (70% genes) but only 54% of isoforms produced unique open-reading frames (ORFs). We found that males exhibited greater isoform richness than females in both tail and gonad tissues but produced a smaller proportion of isoforms with unique ORFs, suggesting that much of the increased isoform variation is unlikely to expand proteomic complexity and may instead reflect stochasticity during splicing rather than intentional transcriptional products intended for translation. Despite widespread AS, we found only 2.3% of genes exhibited sex-biased isoform switching, and only 52% of these switches generated distinct sex-biased ORFs. Together, our long-read data suggest that although isoform diversity is more extensive than previously appreciated, most alternative isoforms are unlikely to generate novel proteins. Instead, a relatively small number of sex-biased isoforms may disproportionately contribute to proteomic divergence between the sexes.

## Introduction

Alternative splicing (AS) is a regulatory mechanism that generates multiple transcript structural variants (isoforms) from a single gene through variable splice site selection. Following the discovery of the split structure of eukaryotic genes, AS was immediately proposed as a mechanism for expanding proteomic complexity without gene duplication (Gilbert 1978; Wright et al. 2022). AS has subsequently been implicated in diverse biological processes, including organ development (Mazin et al. 2021), cell type differentiation (Fiszbein and Kornblihtt 2017) and sex determination (Burtis and Baker 1989), among other physiological and cellular functions (reviewed in Marasco and Kornblihtt 2023; Nilsen and Graveley 2010; Kalsotra and Cooper 2011; Verta and Jacobs 2022). However, there is mounting evidence that many isoforms are non-functional byproducts of splicing stochasticity and other neutral processes (Pickrell et al. 2010; Saudemont et al. 2017; Wan and Larson 2018; Bénitière et al. 2024; Fair et al. 2024; Mi et al. 2026). Consequently, the extent to which observed isoform diversity reflects meaningful increases in proteomic, and therefore phenotypic, complexity remains an open question.

A major limitation in the study of AS has been the difficulty of reconstructing complete isoform structure from short-read RNA sequencing, because individual reads are too short to determine which distant exons occur together within the same transcript (Steijger et al. 2013). As a result, short-read approaches typically quantify local splicing events using reads that directly map across splice junctions of adjacent exons. These approaches are highly conservative and are thus likely to underestimate both isoform richness and biologically meaningful differences in splicing. Long-read RNA sequencing overcomes this limitation by directly sequencing full-length transcripts, allowing isoforms to be identified without bioinformatic reconstruction (Monzó et al. 2025). Consistent with this expectation, a systematic comparison of long-read transcriptomes across multiple datasets from human, mouse and manatee revealed substantially greater isoform richness than previously appreciated (Pardo-Palacios et al. 2024). However, it remains unclear whether the expanded isoform diversity captured by long-reads reflects previously undetected and biologically meaningful AS or simply captures additional transcriptional variation and splicing noise (Su et al. 2024). Consequently, an important unresolved question is the extent to which long-read sequencing provides a more complete picture of biologically relevant splicing differences than short-read approaches.

Long-read sequencing also provides an opportunity to evaluate the functional consequences of AS. If isoform diversity contributes to proteomic complexity, then increases in isoform diversity should frequently be accompanied by increases in the number of distinct open-reading frames (ORFs). However, AS can also alter untranslated regions (Aspden et al. 2023), result in a loss of coding (incomplete ORF), or produce transcripts that are unlikely to generate novel proteins by introducing premature stop codons or retaining introns. These isoforms may instead influence gene regulation or represent selectively neutral variation (Braunschweig et al. 2014; Kurosaki et al. 2019). Determining the relative prevalence of these different outcomes is critical to evaluating the functional potential and evolutionary significance of isoform diversity.

These questions are particularly relevant in the context of sexual dimorphism. Males and females share nearly identical genomes yet often exhibit striking phenotypic differences, making regulatory variation a central mechanism underlying sex-specific trait expression (Mank 2017). Although sex-biased gene expression has been extensively studied for its role in the regulatory divergence of the sexes (Ellegren and Parsch 2007; Mank 2017), much less is known about the role of AS, despite growing evidence that sex-biased splicing is widespread (Telonis-Scott et al. 2009; Gibilisco et al. 2016; Naftaly et al. 2021; Rogers et al. 2021; Singh and Agrawal 2023; Guo et al. 2024; Darolti et al. 2026). If AS contributes to sexual dimorphism through proteomic diversification, sex-biased splicing should frequently generate distinct isoform ORFs or alter protein-coding potential in a sex-specific manner. Alternatively, sex-biased splicing may include a substantial fraction of non-coding or lowly expressed isoforms (Elliott and Grellscheid 2006; Soumillon et al. 2013) that do not contribute meaningfully to phenotypic diversity. Quantifying the contribution of each of these possibilities is essential for understanding how AS contributes to the molecular basis of sexual dimorphism.

To determine the extent to which AS may facilitate transcriptional differentiation of males and females, we compared matched long-read (PacBio Iso-Seq) and short-read (Illumina RNA-seq) datasets generated from identical male and female tail (representing somatic tissue) and gonad (representing reproductive tissue) samples of the guppy (*Poecilia reticulata*), a well-established model for the evolution of sexual dimorphism (Bisazza 1993; Magurran and Garcia 2000; Sharma et al. 2014; van der Bijl et al. 2025). By first comparing the number and overlap of isoforms and splicing events detected by short and long-read RNA-seq, we show that the full-length reads from long-read sequencing reveal substantially more isoforms and more sex differences in isoform usage than event-based analyses from either short or long-read data. Using our long-read dataset, we found that though males have greater isoform richness in both somatic and gonadal tissue compared to females, this increased male isoform richness does not result in substantially increased proteomic complexity, suggesting much of it may be non-functional. Supportive of this, we found that despite widespread sex differences in splicing, only a small subset of genes showed evidence of dynamic isoform switching between the sexes resulting in sex-biased ORFs. This suggests that though AS plays a role in the proteomic divergence of the sexes, much of the increased isoform diversity we observe does not result in increased proteomic complexity.

## Results

### Improved isoform detection with long-reads and isoform-level analysis

We compared genes expressed in both long and short-read datasets (n=11,506) and found 2.7 times more isoforms with long-reads than short-reads (56,957:20,843 Figure 1A and 1B). Only 17% of isoforms were detected in both datasets despite concordance in gene expression quantification (Figure 1C). 68.7% of isoforms were unique to long-reads, consistent with recent studies highlighting the substantially greater isoform complexity captured by long-read sequencing (ex. Li et al. 2018; Naftaly et al. 2021). A smaller fraction of isoforms (14.3%) was unique to the short-read assembly. To assess whether these represent short-read assembly artefacts or low-abundance isoforms that may have simply been removed from the long-read assembly during filtering, we examined their splice-junction composition across datasets. Of these, 39.5% contained junctions also observed in the long-read dataset but assembled into different isoforms. We found the other 60.5% included at least one junction not detected in the filtered long-read dataset and that isoforms containing novel junctions had lower expression than both shared-junction and shared-isoform categories (Figure S1). Given that all junctions in our long-read assembly are cross-validated with short-reads (see Methods) novel-junction isoforms are more likely to reflect low abundance transcripts removed during long-read filtering, whereas shared-junction isoforms are more consistent with putative short-read assembly errors, though these explanations are not mutually exclusive.

Focusing on splicing events (Figure 1D), we found that only 6.9% of all events were shared between datasets and that the number of events unique to long-reads was 7x more than those unique to the short-read assembly (59,148:7,613; Table S1). We identified more events across all event types with long-reads, including those considered most likely to contribute to proteome expansion (exon skipping (ES) and mutually exclusive exons (MX; Weatheritt et al. 2016) as well as event types with less clear functional purpose (intron retention (IR) and alternative 3’ and 5’ splice sites (A3/A5)).

A3/A5 events were by far the most abundant in the long-read dataset with over 3x more events detected than the second most common event type (IR; Figure 1E). We found these two categories had the lowest overlap between datasets with only 6.3% of A3/A5 events and 1.4% of IR events shared. Alternatively, ES was the most abundant event type recovered in the short-read dataset with the largest overlap between datasets (15.3%).

We next tested for sex-biased splicing by calculating the difference in percent-spliced-in (ΔPSI) between the sexes for shared events. We identified nearly 3x more significant events with long-reads with little overlap again (Figure 1F). The degree of concordance differed across event types with ES events showing the greatest overlap (13.4%) and IR events showing the least (<1%). We found the greatest number of sex-biased splicing events were A3/A5 events in the long-read dataset followed by ES events, but this order was reversed for short-reads. The least common in both datasets were MX events which were also the least frequently detected event type overall. For those events that were detected in both conditions, ΔPSI values were positively correlated (Pearson correlation, *r* = 0.46, p < 0.001; Figure S2).

We next asked whether isoform-level analysis could reveal additional sex-biased splicing that was missed by event-level approaches. Using our long-read dataset, we tested for differential isoform usage (DIU) between males and females, identifying genes in which individual isoforms showed sex-dependent changes in abundance relative to the overall change in gene expression. Isoform-level analysis revealed a strikingly greater extent of sex-biased splicing than event-level analyses with 38.4% of genes expressed in the gonad and 0.8% of genes expressed in the tail showed sex-biased isoform usage. In the gonad, this represented a 3.2x increase over long-read event-level analysis (12.0%) and a 6.6x increase over short-read event-level analysis (5.8%; Table 1). The difference was more modest in the tail, where isoform-level analysis identified 0.8% of genes compared with 0.3% and 0.4% using long and short-read event-level analyses respectively. Together, these results suggest that both the data used for transcriptome assembly and quantification (short or long-reads) and the level at which AS is detected (event or isoform-level) can have a profound effect on the number of sex-biased splicing genes detected, with isoform-level analysis revealing substantially more sex-biased regulation than event-level approaches, particularly in the gonad.

Using this improved estimate of sex-biased splicing, we examined the overlap with genes that were differentially expressed at the gene level. In the tail, 11.5% of sex-biased splicing genes were also differentially expressed, compared with 58.5% in the gonad (Figure S3), suggesting these two regulatory mechanisms often co-occur to produce a complex regulatory architecture, especially in reproductive tissue.

**Figure 1:**
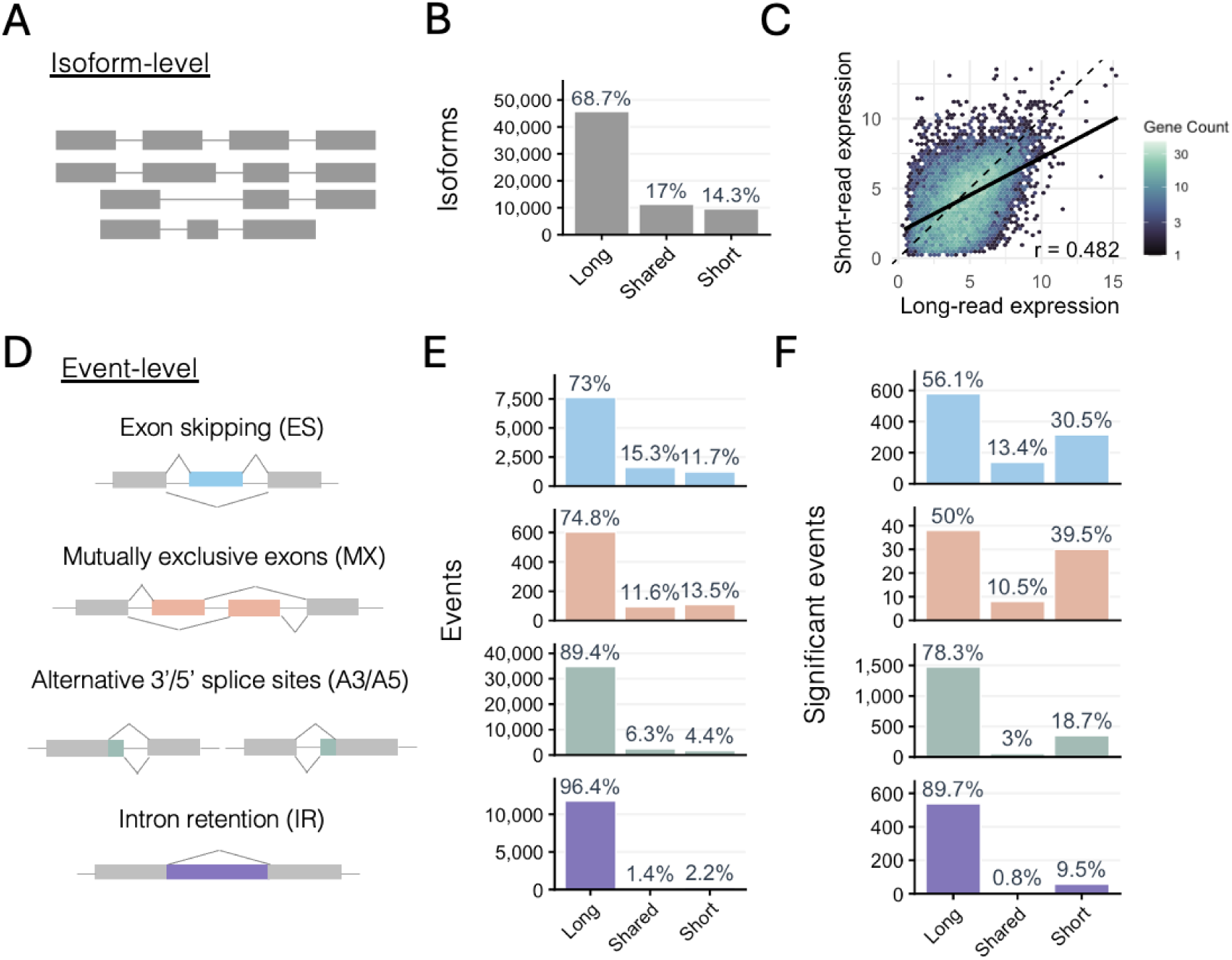
(A) Alternative splicing (AS) measured at the isoform-level using full-length assembled transcripts (B) Barplot illustrating the proportion of assembled isoforms detected by each dataset. (C) Comparison of gene expression (log_2_(TPM + 1)) as measured with long-reads against the long-read transcriptome and short-reads against the short-read transcriptome. Pearson correlation *r* = 0.482, p < 0.001. Hexagons represent genes and are coloured by overlapping genes following the legend. Dotted black line shows 1:1 line, solid black line shows regression line. (D) AS can be measured locally at the event-level using reads spanning splice junctions. For each event type constitutive exons are in light grey and AS patterns are mapped via paths above and below each pre-mRNA. AS exons are coloured by event type (blue: exon skipping, ES; orange: mutually exclusive exons, MX; green: alternative 3’ or 5’ splice sites, A3/A5; purple: intron retention, IR). (E) Barplots illustrating the proportion of event detected by each dataset faceted by event type. Colours follow those outlined in D. (F) Barplots as in E but showing only the proportion of shared events which were found to be alternatively spliced between the sexes (ΔPSI > 0.2, FDR < 0.05).

**Table 1:** Number of genes exhibiting sex-biased splicing depending on the read type and AS detection method used. For event-level analyses, only genes expressed in both sexes and in both the short- and long-read datasets were included, allowing direct comparison of sequencing technologies. For isoform-level analysis, only genes expressed in both sexes were included. The table illustrates the substantially greater number of sex-biased splicing genes detected by isoform-level analysis compared with event-level approaches, particularly in gonad tissue.

| Tissue | Read type | AS detection method | Genes tested | Genes with sex-biased splicing | Percentage of genes with sex-biased splicing |
| --- | --- | --- | --- | --- | --- |
| Tail | Short | Event-level | 10,395 | 37 | 0.4% |
| Tail | Long | Event-level | 10,395 | 34 | 0.3% |
| Tail | Long | Isoform-level | 12,675 | 96 | 0.8% |
| Gonad | Short | Event-level | 10,999 | 634 | 5.8% |
| Gonad | Long | Event-level | 10,999 | 1,325 | 12.0% |
| Gonad | Long | Isoform-level | 13,138 | 5,042 | 38.4% |
| All | Short | Event-level | 11,506 | 657 | 5.7% |
| All | Long | Event-level | 11,506 | 1,246 | 10.8% |
| All | Long | Isoform-level | 13,721 | 5,060 | 36.9% |

### Alternative splicing is common, but many isoforms do not produce unique ORFs

AS was widespread in our long-read dataset with a mean isoform richness (Figure 2A) of 4.67 [4.56, 4.78] per gene (mean [95% CI, mean ± 1.96 standard errors]). We found 69.8% of genes expressed at least two isoforms, while 30.3% expressed five or more (Figure 2B). Across all protein-coding genes only 54% of isoforms produced unique ORFs, with the next largest category being isoforms with retained introns (18.2%; Table S2). Both gene and isoform expression clustered by tissue first and then sex, with little differentiation between the sexes in tail samples (Figure S4).

Isoform expression was often highly skewed within a gene (Figure 2C). To quantify this skew, we calculated isoform diversity, defined as the effective number of equally expressed isoforms corresponding to the observed expression distribution. Mean isoform diversity across all samples was 1.99 [1.96, 2.02], indicating that gene expression was typically distributed among isoforms in a manner equivalent to approximately two equally expressed isoforms, despite the greater number of isoforms detected per gene. We also found that isoform diversity was not related to overall gene expression (linear regression, R^2^ < 0.01, p < 0.001; Figure S5). We further classified the most highly expressed isoforms as effective based on our measure of diversity (Figure 2A and Methods). We found ineffective isoforms to not only have lower isoform expression but also to more frequently have incomplete or absent ORFs (i.e. they are non-coding), retained introns, or premature stop-codons, all of which suggest that lowly expressed isoforms tend to have lower translation potential (Figure S6).

Consistent with this, we observed that though isoform richness correlated with total gene expression (linear regression, R^2^ = 0.05, p < 0.001; Figure S5), the proportion of isoforms with unique ORFs decreased as isoform richness increased, such that genes with higher richness express more isoforms with redundant ORFs or those putatively targeted for nonsense-mediated decay (NMD) via intron retention or premature stop codons (Braunschweig et al. 2014); Figure 2D and 2E).

**Figure 2:**
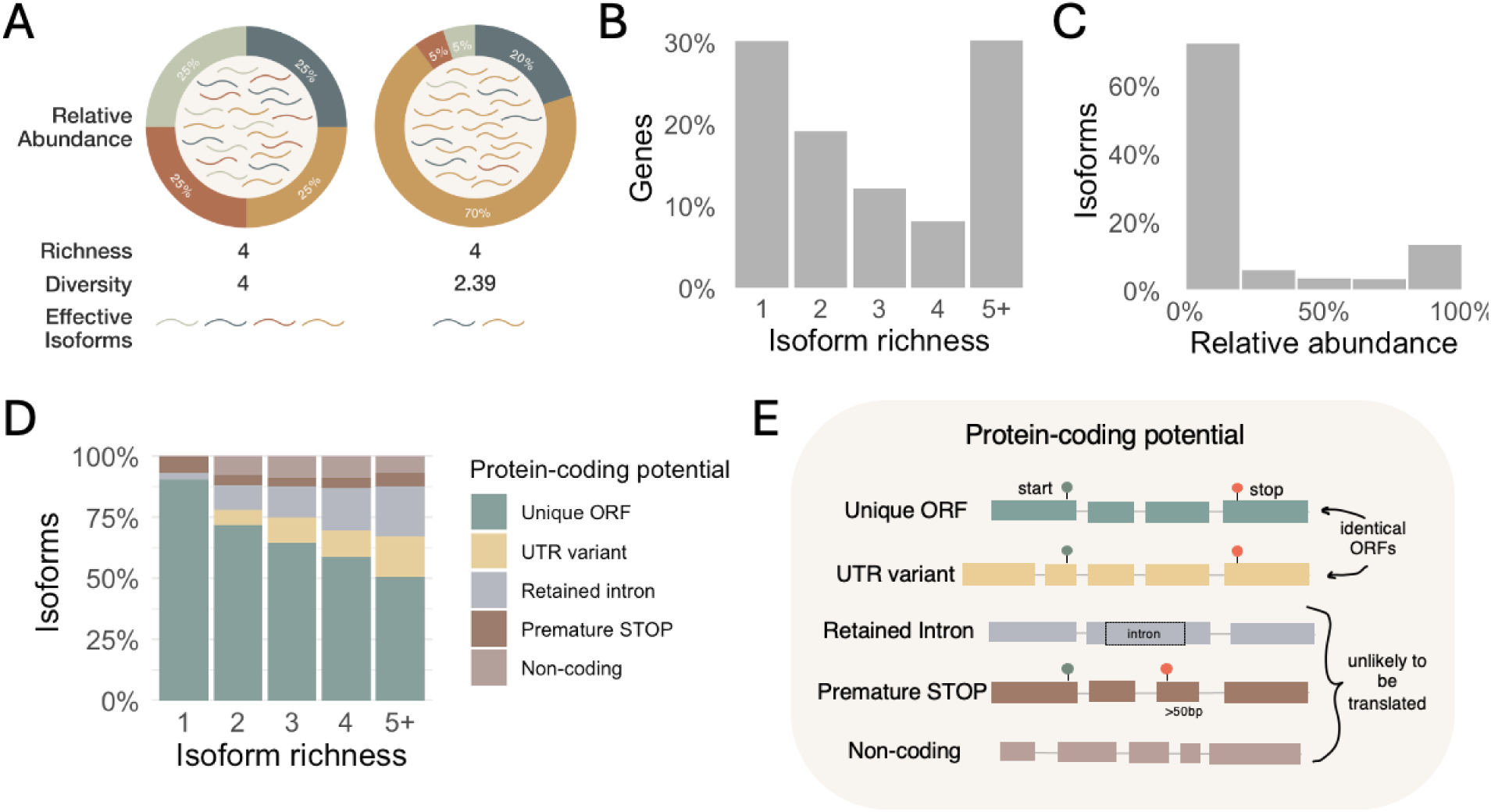
(A) Schematic demonstrating the difference between isoform metrics used in this study where the donut plots show the relative abundance (proportion of total gene expression contributed by a single isoform) of the four distinct isoforms each denoted with a different colour. Isoform richness, which is simply the raw count of unique isoforms per gene, is the same for both examples, whereas isoform diversity differs because it accounts for the evenness of expression. We quantified isoform diversity as 2*^H^*,where *H* is the Shannon entropy of isoform expression within a gene, calculated as *H* = − ∑ *p_i_*ln *p_i_* where *p_i_* is the relative abundance of isoform *i*. Thus, isoform diversity represents the effective number of equally expressed isoforms corresponding to the observed expression distribution. We then labelled isoforms as either effective or ineffective by rounding isoform diversity to the nearest whole number (*n*) and selecting the *n* most highly expressed isoforms as effective and all remaining isoforms as ineffective (Schertzer et al. 2025). (B) Histogram of isoform richness per gene calculated with the long-read dataset. (C) Histogram of isoform relative abundance. (D) Stacked barplot showing isoform protein-coding potential across genes binned by the total number of isoforms expressed (isoform richness). (E) Schematic demonstrating how isoforms were categorized by protein-coding potential using both their exon structure and ORF prediction. **Unique ORF**: an isoform with a complete ORF observed for the first time in the dataset; **UTR variant**: an isoform with a complete ORF that has previously been observed in the dataset and therefore differs only in UTR structure; **retained intron**: an isoform containing a retained intron and therefore unlikely to be translated due to either nonsense-mediated decay or nuclear retention (Jacob and Smith 2017); **premature STOP**: an isoform containing a premature stop codon (located >50bp upstream from the last exon-exon boundary) and therefore likely subject to nonsense-mediated decay (Kurosaki et al. 2019); **non-coding**: an isoform lacking a complete ORF, as determined by *TD2* (Mao et al. 2025).

### Males have higher isoform richness and diversity in both somatic and germline tissues

We observed elevated transcriptome complexity in males across replicates, with males showing higher isoform richness and diversity than females in both tissues (Figure 3). To quantify whether this was a consistent genome-wide effect we calculated the male:female log rate ratio of isoform richness for each gene (Figure 3A) and found that males produced 9.6% [9.0%, 10.3%] (mean log effect [95% CI, mean log effect ± 1.96 standard errors]) more isoforms than females in the tail and 5.5% [4.3%, 6.8%] more in the gonad. The same was true for isoform diversity (Figure 3D), males produced 3.6% [3.3%, 3.9%] more effective isoforms in the tail and 4.2% [3.5%, 4.9%] more in the gonad.

We identified substantial sex differences in gonad isoform composition, with significant shifts across multiple isoform types (Figure 3C; Table S2). Male gonads showed a reduction in unique ORF isoforms and an increased proportions of isoforms in non-coding, premature STOP, and UTR variant categories. These results indicate a restructuring of isoform composition in male gonads toward isoforms with less unique protein-coding potential. Sex differences in tail tissue were considerably weaker, with significant changes limited to unique ORF and UTR variant categories. We repeated this analysis with only effective isoforms and found the pattern held in gonads but not in tails (Figure 3F).

**Figure 3:**
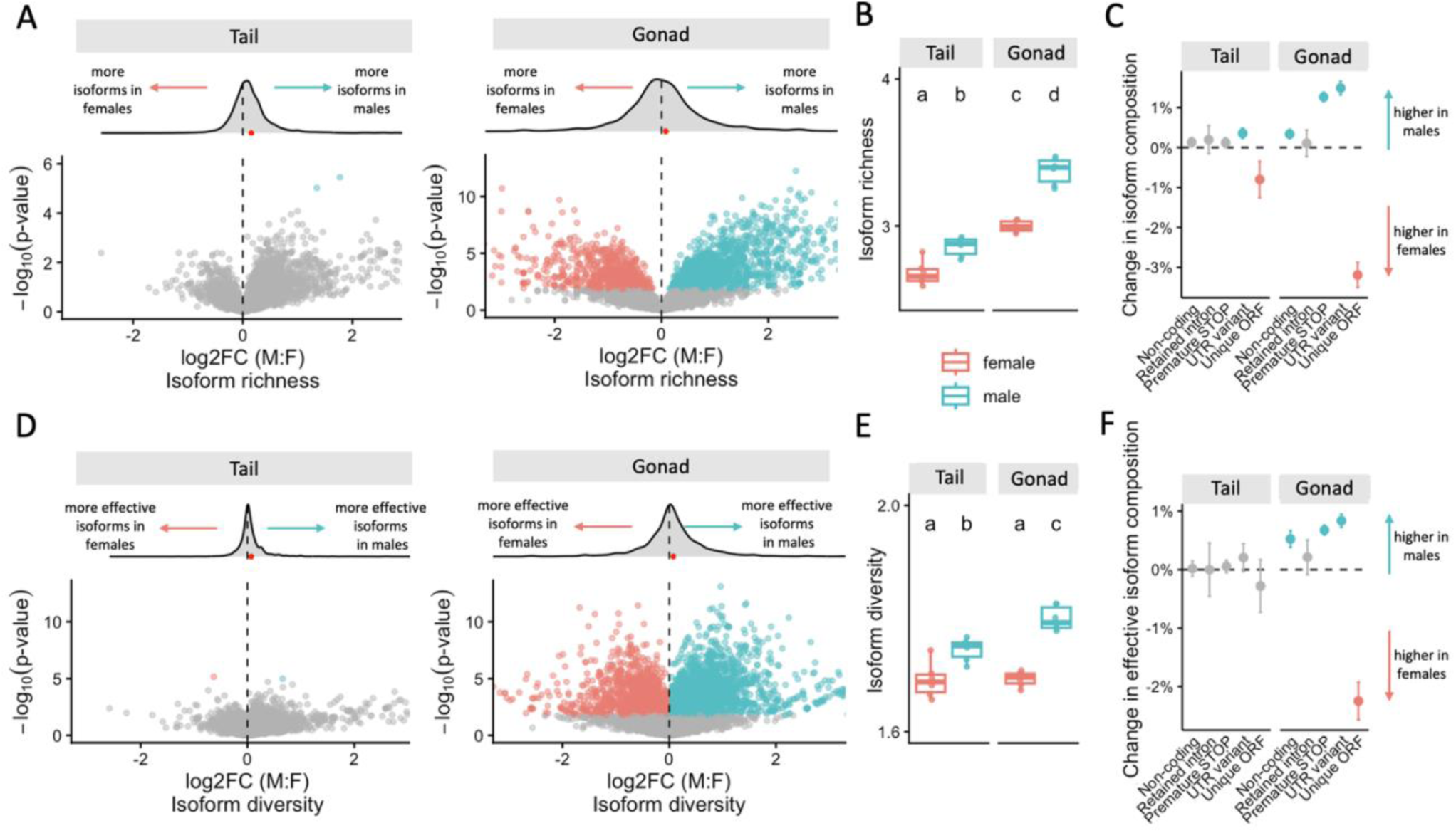
(A) Volcano plots showing the log_2_ male:female fold change in isoform richness for genes expressed in both sexes. Points are coloured by statistical significance (FDR < 0.05), with significant genes coloured according to the sex exhibiting greater isoform richness (0 > log_2_FC, female, red; 0 < log_2_FC, male, blue). X-axes are cropped to capture core of distributions. (B) Boxplot of isoform richness for each sex for genes expressed in both sexes. Boxes represent interquartile range and points represent the mean of each biological replicate. Letters above boxplots indicate statistically significant differences among groups based on Kruskal-Wallis tests followed by Dunn’s post hoc pairwise comparisons with Benjamini-Hochberg correction for multiple testing (FDR < 0.05). (C) Forest plot showing mean differences in isoform composition between males and females (male−female) estimated via generalized linear models fit within each tissue. Proportions were calculated per sample across all isoforms from genes expressed in both sexes. Error bars represent 95% confidence intervals (mean difference ± 1.96 standard errors). Colours indicate effect direction and statistical significance (FDR < 0.05). Isoform composition comprises five structural categories based on protein-coding potential (Fig 2D; Methods). (D-F) Equivalent analyses to panels A–C, but for isoform diversity and effective isoforms, respectively.

### Male-biased genes have the highest isoform diversity

We next used the sex-bias categories defined in the previous differential gene expression analysis to examine whether isoform richness and diversity differed among sex-biased and unbiased genes. Given the much greater number of sex-biased genes detected in the gonad, we focused these comparisons on gonadal genes only (Figure S3; Table S3). We observed a consistent relationship between isoform metrics and sex-biased expression in the gonad, with male-biased genes showing the highest isoform richness and diversity in both sexes (Figure 4) while female-biased genes were the least diverse in both sexes. For isoform richness, differences between the bias categories were primarily driven by changes in males, suggesting a relationship with male expression level (Figure 4A and 4B).

Analysis of isoform composition revealed pronounced differences between male and female-biased genes in the gonad (Figure 4C; Table S4). Male-biased genes showed a significant shift away from isoforms producing unique ORFs, the category associated with increased proteomic diversity. Instead, male-biased genes produced more isoforms that were non-coding, had premature stop codons, retained introns, or varied only in the UTRs. These effects were strongest in male samples, but the same overall pattern was observed in female samples. We repeated this analysis after filtering for only effective isoforms and found the result held in both sexes (Figure 4F).

To investigate why male-biased genes had greater isoform richness and diversity, we asked whether these patterns could be explained by gene expression level and the opportunity for stochastic mis-splicing. Previous studies have shown that isoform richness can increase with gene expression because greater transcriptional activity provides more opportunities for splicing errors (Pickrell et al. 2010; Bénitière et al. 2024). Under this assumption, the strength and direction of the relationship between expression and isoform richness provides an indication of the extent to which additional isoforms may arise from transcriptional noise, where a stronger positive relationship is consistent with a greater contribution of stochastic mis-splicing. We therefore examined the relationship between gene expression and isoform richness separately for each condition and found this relationship was strongly sex- and tissue-specific. In the male gonad, gene expression explained 10% of the variation in isoform richness, compared with only 1–2% in all other conditions (Figure S7, p < 0.001), consistent with a greater contribution of transcriptional noise to isoform production in the male gonad.

We next looked at isoform diversity which captures how even isoform expression is within a gene. A negative relationship between gene expression and isoform diversity is therefore consistent with stronger constraint on isoform usage as expression increases, whereas the absence of such a relationship suggests weaker constraint on splicing fidelity. We found that isoform diversity was unrelated to gene expression in the male gonad, whereas all other conditions showed a negative relationship (R² = 0.03, p < 0.001; Figure S7). Taken together, the stronger and more positive relationship between gene expression and isoform richness and the absence of a relationship between gene expression and isoform diversity distinguish the male gonad from the other conditions and suggest reduced constraint on splicing fidelity in males.

To further test whether this pattern was consistent with stochastic isoform production, we compared our observations with a null model in which additional isoforms arise from opportunities for mis-splicing and a baseline rate of mis-splicing (0.7% per intron; Pickrell et al. 2010; see Methods). The model assumes that genes intend to produce a single isoform and that additional isoforms arise from stochastic splicing errors, with gene expression level and intron number determining the number of interactions with the spliceosome and thus opportunities for such errors. This model therefore provides a null expectation for isoform richness and diversity in the absence of sex or tissue-specific differences in splicing fidelity. Across tissues, the model generally overestimated isoform richness and diversity among highly expressed genes, indicating that observed isoform production was more constrained than expected from stochastic mis-splicing alone (Figure S7). In contrast, the male gonad showed the closest agreement between observed and simulated values. Thus, among the conditions examined, the male gonad most closely resembled the expectation under stochastic isoform production. This supports our interpretation that the excess isoforms observed in males may arise in part from transcriptional noise rather than increased production of functionally regulated isoforms.

**Figure 4:**
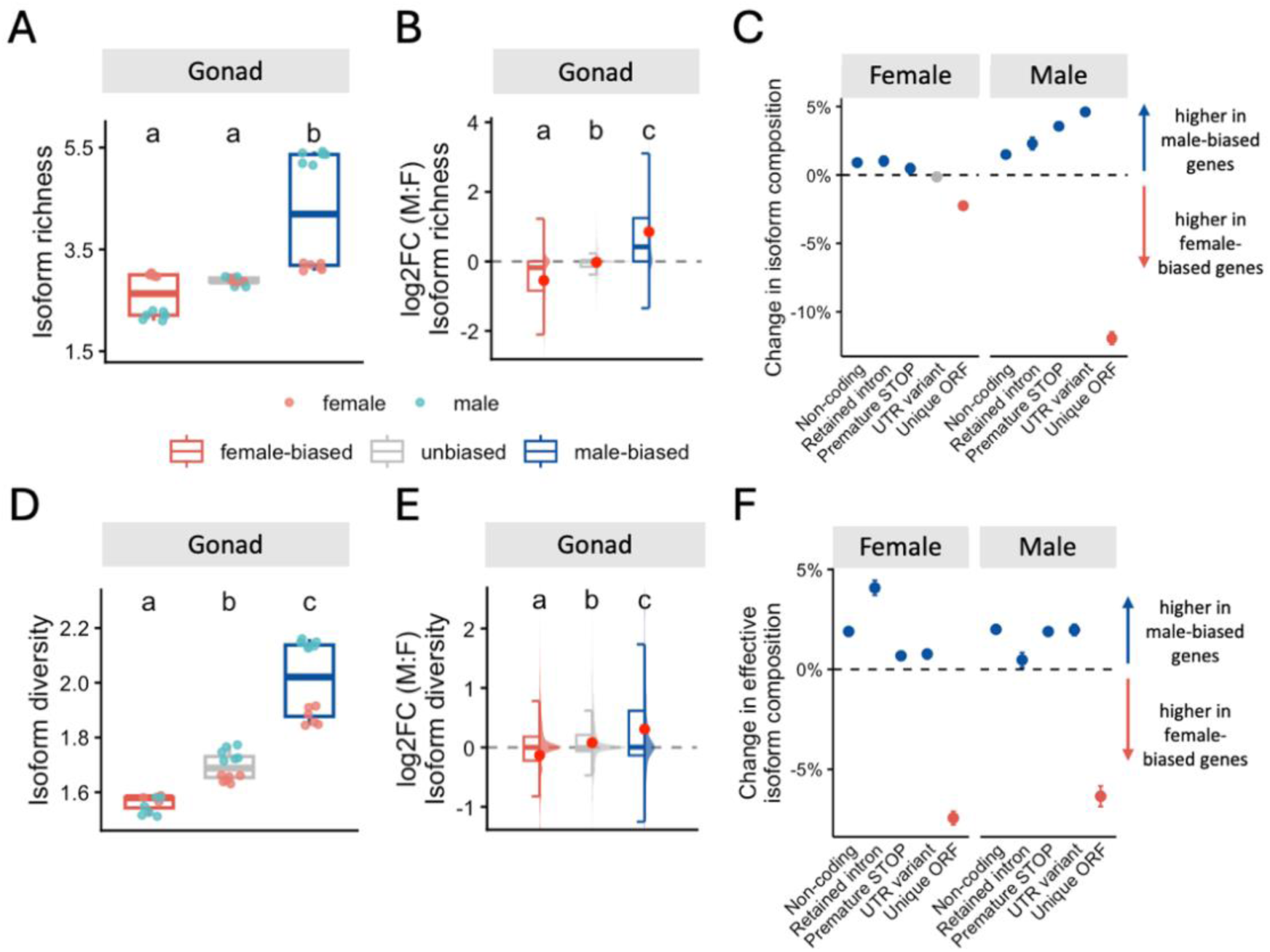
(A) Boxplot of gonad isoform richness for genes with female-biased, unbiased and male-biased gene expression. Female-biased: log_2_FC < -1, unbiased: -1 <= log_2_FC <= 1, male-biased: 1 < log_2_FC. Boxes represent interquartile range and points represent the mean of each biological replicate. (B) Male:female fold change shown for isoform richness of gonad genes with female-biased, unbiased and male-biased expression. Y-axis is cropped to show core of distribution. (C) Forest plot showing mean differences in isoform composition between sex-biased genes in the gonad (male-biased−female-biased) estimated via generalized linear models fit within each sex. Proportions were calculated per sample across all isoforms from genes expressed in both sexes. Error bars represent 95% confidence intervals (mean difference ± 1.96 standard errors). Colours indicate effect direction and statistical significance (FDR < 0.05). Isoform composition comprises five categories based on protein-coding capacity. (D-F) The same as panels A-C but for isoform diversity and effective isoforms respectively. Letters above all boxplots indicate statistically significant differences among groups based on Kruskal-Wallis tests followed by Dunn’s post-hoc pairwise comparisons with Benjamini-Hochberg correction for multiple testing (FDR < 0.05).

### Putative evidence for isoform switching including complex multi-isoform events

We next quantified differential gene expression (DGE) and differential isoform expression (DIE) between males and females using absolute expression levels. Sex-biased expression was rare in the tail, affecting only 1.6% of genes and 0.6% of isoforms, but was widespread in the gonad, where 54.8% of genes and 44% of isoforms exhibited sex-biased expression (Table S3). Gene-level bias did not consistently extend to all constituent isoforms, with gene and isoform-level bias directions exhibiting only moderate agreement in both tail (Cohen’s unweighted κ = 0.393, p < 0.001) and gonad tissues (Cohen’s unweighted κ = 0.493, p < 0.001). In tail tissue, we found 78.1% of isoforms from female-biased genes and 72.1% of isoforms from male-biased genes exhibited unbiased isoform expression. Similarly, in gonad tissue, we found that 38.0% of isoforms produced by female-biased genes and 38.6% of isoforms from male-biased genes showed unbiased isoform expression. Conversely, unbiased genes produced sex-biased isoforms only 0.20% of the time in the tail (0.02% female-biased, 0.18% male-biased) but 20.0% of the time in the gonad (10.1% female-biased, 9.9% male-biased). These results demonstrate that gene-level analyses only partially capture the full transcriptional complexity observable at the isoform-level.

We next asked whether AS provides a mechanism for sex-specific proteomic expansion through isoform switching. Specifically, we looked for cases where males and females preferentially expressed different isoforms of the same gene (see Methods). We identified 319 genes exhibiting isoform switching in the gonad, but none in the tail, indicating that this phenomenon is largely restricted to the reproductive tissue. Strikingly, these events did not always conform to the classic model of a simple 1:1 isoform switch, in which one female-biased isoform in females is replaced with one male-biased isoform in males. Instead, isoform switching was frequently far more complex, with individual genes often producing multiple sex-biased isoforms within one or both sexes (Figure 5A), revealing extensive transcriptome remodeling rather than straightforward isoform replacement.

We further investigated whether isoform switching genes also retained unbiased isoforms expressed in the same tissue, which could indicate retention of ancestral functions and potential sub-functionalization. We observed this pattern in 26% of isoform-switching genes (83/319; Figure 5B).

Across all isoform switching types, the majority of events resulted in the production of sex-biased ORFs, with distinct coding repertoires between males and females (52%; 166/319). In contrast, a subset of switches involved changes in protein-coding potential (Table S5), including cases where coding capacity was lost in one sex. Among these events, loss of coding occurred more frequently in males than females (48/72; 66.7%, 95% CI [54.6%, 77.3%]), representing a significant deviation from a 50:50 expectation (exact binomial test, p = 0.006; Figure 5C).

**Figure 5:**
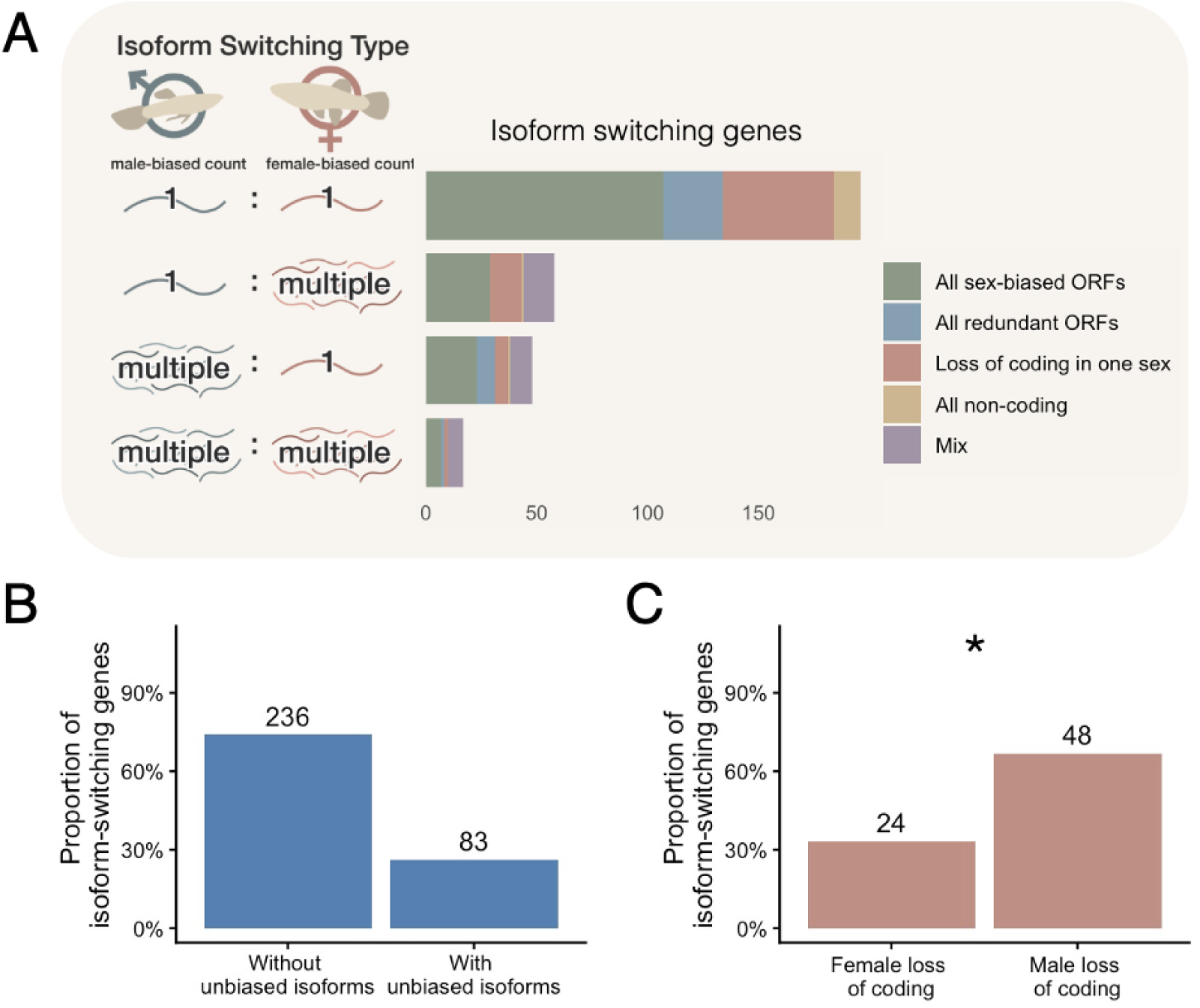
(A) Histogram of isoform switching genes categorized by the number of sex-biased isoforms they produce in males and females. Bars are coloured to show the relative proportion of each protein-coding outcome. **All sex-biased ORFs**: both males and females produce distinct, sex-biased ORFs; **all redundant ORFs**: both males and females produce the same ORF but diverge in their exon structure in the UTRs; **loss of coding in one sex**: all sex-biased isoforms expressed in one sex are classified as either non-coding, premature STOP or intron retention, while the other sex retains protein-coding isoforms; **all non-coding**: all sex-biased isoforms expressed in both sexes lack protein-coding isoforms (unique ORF or UTR variant categories); **mix**: complex switching events that do not fit cleanly into the above categories. (B) Proportion of isoform-switching genes with and without unbiased isoforms expressed at >5 TPM in both sexes, suggestive of sub-functionalization. (C) Proportion of isoform-switching genes with loss of coding in males versus females. Asterisk denotes significant deviation from binomial expectation (exact binomial test, p = 0.006).

## Discussion

### Long-reads improve detection of sex-biased splicing

Consistent with recent long-read evaluations (Monzó et al. 2025; Pardo-Palacios et al. 2024; Schertzer et al. 2025), we identified a 2.7-fold increase in the number of isoforms detected with long-reads, alongside a 7-fold increase in detected local splicing events. The remarkably narrow overlap between long and short-read approaches, 17% at the isoform-level and 6.9% at the event-level, underscores the structural limitations of isoform assembly using short-reads. Concordant with this, we found that nearly 40% of isoforms that were unique to the short-read dataset showed putative evidence of mis-assembly.

At the event level, the discrepancy in detected events between platforms was largest for alternative 3′ and 5′ splice site (A3/A5) events and intron retention (IR) events which were the most common event types overall. We further found that only 5.8% of significant sex-biased events overlapped between datasets with the disparity between platforms being largest again for A3/A5 and IR events. This is counter to previous short-read studies which have reported that exon skipping is the most common AS event associated with sex differences (Karlebach et al. 2020; Rogers et al. 2021; Lu et al. 2022). Together this indicates that biological conclusions regarding sex-biased AS are heavily contingent upon the sequencing technology utilized and may bias our understanding of splicing towards outcomes associated with increased proteomic diversity.

Overall, long-read sequencing coupled with isoform-level analyses substantially expanded the landscape of detected sex-biased splicing (Table 1 and 2). While event-level analyses identified sex-biased splicing in only 5.7% of genes using short-reads and 10.8% of genes using long-reads, long-read isoform-level analyses increased this to 36.9% of genes. This expanded view of isoform regulation further revealed that sex-biased splicing frequently co-occurs with differential gene expression, with 58% of sex-biased splicing genes also showing DGE. Thus, rather than representing mutually exclusive regulatory strategies as previously proposed (Jacobs and Elmer 2021; Singh and Agrawal 2023), AS and gene expression appear to act together to generate a complex regulatory landscape underlying sex-specific transcriptional variation.

Overall, the discrepancy between isoform and event-level approaches, even when using the same long-read data, highlights that many biologically relevant splicing changes cannot be captured by the discrete splicing event categories typically screened for. Instead, sex-biased AS often involves complex patterns of coordinated exon and intron usage changes co-occurring with changes in expression level which are obscured by event-centric approaches.

**Table 2:**
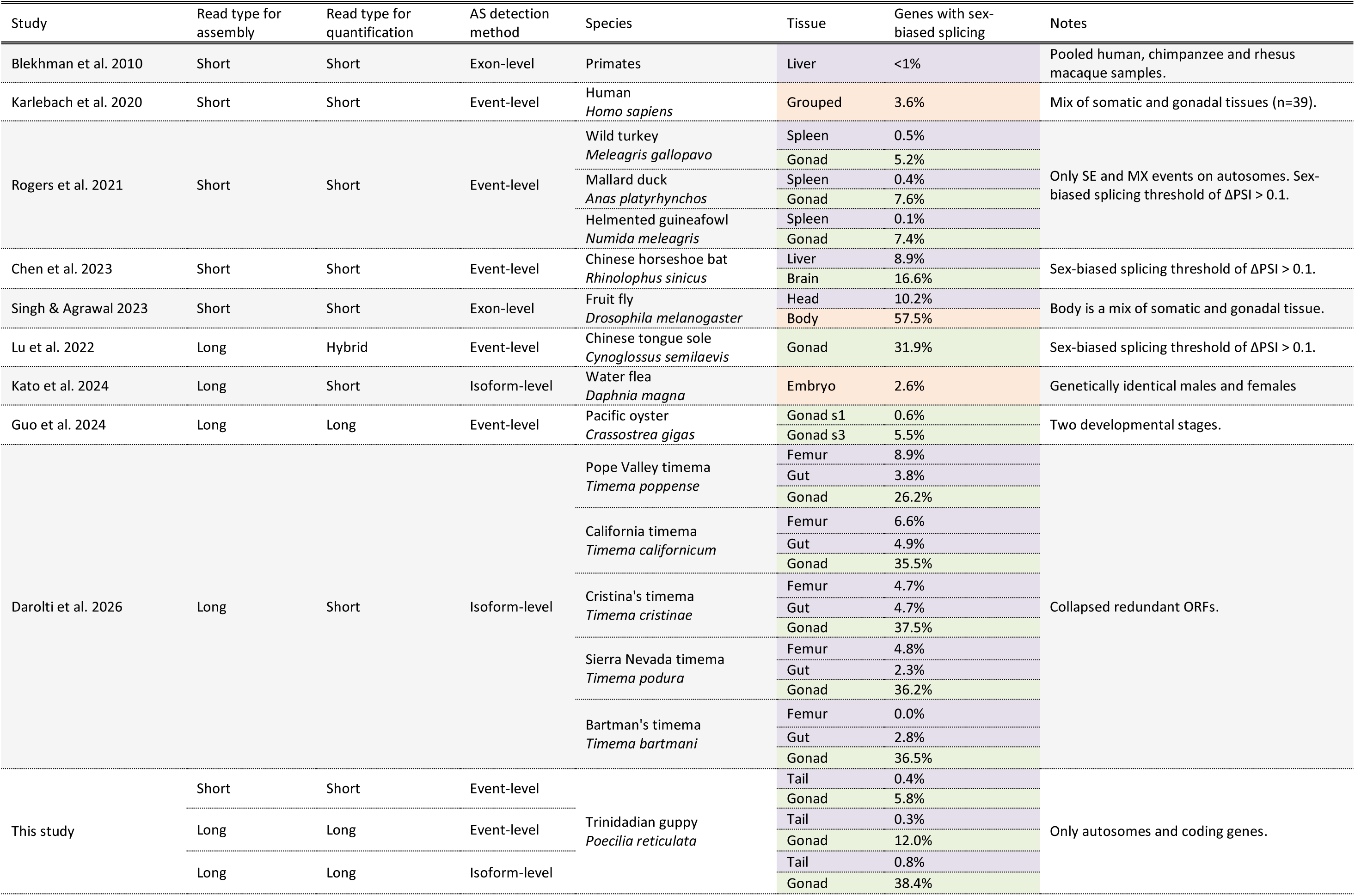
Comparison of the proportion of sex-biased splicing genes found in recent animal studies showing the sequencing technology used for transcriptome assembly and expression quantification, and the AS detection method used. Long-read transcriptome assembly paired with isoform-level AS detection consistently detects more sex-biased splicing, especially in the reproductive tissue. Reproductive tissues are coloured in green, somatic tissues in purple and mixed/bulk tissues in orange.

### Diminishing returns on isoform complexity

Although AS is undeniably widespread in the guppy, with nearly 70% of expressed genes yielding multiple isoforms, we found that just under half (46%) of all isoforms did not produce unique ORFs and instead contained retained introns, premature stop codons or varied exclusively in the untranslated regions (UTRs). Further, we found that fewer than half of the detected isoforms are effective contributors to overall gene expression. Instead, isoform expression within a given gene is highly skewed, meaning that rare, low-abundance variants make up the majority of isoform counts but represent negligible fractions of total gene expression level. These ineffective isoforms showed evidence of reduced translation potential, with fewer complete ORFs than found in effective isoforms.

Globally, we observed a positive relationship between gene expression level and isoform richness but no relationship between gene expression level and isoform diversity. This is consistent with previous studies which reported a positive relationship between gene expression level and incidence of mis-splicing, likely due to increased interactions with the spliceosome (Wan and Larson 2018). It also suggests that highly expressed genes experience stronger selection on splicing fidelity (Bénitière et al. 2024; Saudemont et al. 2017) as these genes produce more absolute isoforms overall but the number of effective isoforms remains consistent across expression levels. Together, these results support previous findings suggesting that many isoforms are likely the result of stochasticity during splicing rather than intentional transcriptional products intended for translation (Bénitière et al. 2024).

### Males are more transcriptionally complex

Using our long-read dataset to assess differences in isoform richness and diversity, we uncovered pervasive sexual dimorphism in both gonad and tail tissue. While previous studies have documented higher isoform richness in the testes in mammals (Soumillon et al. 2013; Mazin et al. 2021), fish (Naftaly et al. 2021; Lu et al. 2022), mollusks (Guo et al. 2024), and insects (Darolti et al. 2026; but see Gibilisco et al. 2016), our data demonstrate that guppy males exhibit higher isoform richness and diversity than females in both gonad and somatic tail tissue. In gonads, this pattern was partly driven by male-biased genes, which displayed greater isoform richness and diversity than both unbiased and female-biased genes.

This pattern points to two alternative evolutionary scenarios. First, the increased isoform complexity may represent more adaptive AS within males, as many studies have shown accelerated rates of molecular evolution in male-biased genes (Ellegren and Parsch 2007; Mank 2017) with some providing evidence this is driven by positive selection (Ávila et al. 2015). However, other studies have flagged elevated rates of evolution for male-biased genes as ultimately a product of relaxed constraint (Gershoni and Pietrokovski 2014; Harrison et al. 2015; Tosto et al. 2023). Consistent with the latter case, elevated isoform complexity may arise from increased transcriptional permissiveness or accumulation of neutral splicing variation. For example, open chromatin states during spermatogenesis in mammals can trigger leaky transcription that could lead to the production of more aberrant isoforms (Soumillon et al. 2013).

To distinguish between these possibilities, we examined whether increased isoform richness and diversity were accompanied by shifts in isoform structural composition. Across both tissues, males exhibited reduced proportions of isoforms producing unique ORFs and increased proportions of isoforms varying only in the UTRs. In gonads, males additionally showed increased representation of isoforms that were non-coding or had premature stop codons. These patterns were particularly pronounced among male-biased genes, which produced substantially more alternative isoforms but fewer unique ORFs than female-biased genes. Notably, this compositional shift was observed when male-biased genes were expressed in either sex, indicating that it reflects properties of male-biased genes rather than solely the male transcriptional environment. This enrichment of isoforms that do not expand proteome diversity among male-biased genes suggests that much of the increased isoform output may represent neutral splicing variation rather than adaptive diversification of the proteomic repertoire.

To further test whether increased isoform diversity in males could arise through stochastic mis-splicing, we simulated a null model based on an empirical estimate of a 0.7% mis-splicing rate (Pickrell et al. 2010). While our model consistently overestimated isoform richness and diversity for highly expressed genes across tissues, consistent with selection on splicing fidelity, the male gonad showed the closest concordance with simulated results. Together, these results support the hypothesis that most excess isoforms produced by males are a byproduct of transcriptional noise or relaxed selective constraints rather than expanded adaptive AS in the testis.

### Isoform-level analyses reveal hidden regulatory variation

We found discordance between differential expression measured at the gene (DGE) and isoform (DIE) levels. In the gonad, more than half of all expressed genes exhibit sex-biased gene expression, yet their component isoforms are frequently unbiased. It’s possible that these unbiased isoforms represent noisy mis-splicing that is not phenotypically relevant in either sex, or alternatively these could represent ubiquitous isoforms that are co-expressed with sex-specific isoforms that have undergone sub-functionalization, but further study is required to distinguish these possibilities. These results illustrate how analyses conducted only at the gene level miss much of the molecular architecture that distinguishes the sexes.

### Isoform switching reveals adaptive splicing

The classic example of isoform switching between the sexes involves a binary 1:1 switch between a single male-biased isoform and a single female-biased isoform such as we find in the *Drosophila* sex-determination cascade. This cascade operates through a series of strict, 1:1 splicing switches across genes like *Sex-lethal*, *transforme*r, and *doublesex*, avoiding intermediate or multi-isoform states (Salz and Erickson 2010). This assumption is reinforced by common methods for investigating isoform switching, which focus only on the most highly expressed isoform of each sex. By expanding our search for isoform switching events to any switch found above a certain expression threshold (>5 TPM), we found that isoform switching in guppies often involves more complex dynamics where the sexes deploy multiple sex-biased isoforms concurrently from the same gene. All isoform-switching events were exclusively found in the gonad, consistent with their role in overcoming shared genetic architecture to allow for sex-specific divergence.

In total we found 319 genes that exhibited isoform switching dynamics, representing 2.4% of genes expressed in gonad and 2.3% of genes expressed overall. Our assessment of the structural results of these events indicates that the majority of switching events (52%) produced completely unique ORFs in each sex, demonstrating that whether a gene uses a simple or complex switching strategy, the primary outcome is to generate distinct, sex-specific isoforms from the same underlying sequence. We found that 26% of these events occurred in genes that also produced unbiased isoforms, providing some support for sub-functionalization of alternative sex-biased isoforms.

When switching results in a putative loss of function, male-specific loss of function was nearly twice as common. Notably, there were less complex switching events with multiple male isoforms and only a single female isoform, despite our broader result that males make more isoforms overall. This provides additional evidence that increased isoform diversity in males is not substantially increasing proteomic complexity.

### Concluding remarks

Since its introduction, long-read RNA sequencing has been put forth as a substantial improvement over short-read technologies for isoform detection and splicing inference, but the magnitude of this improvement has yet to be systematically quantified. Our results highlight a profound disconnect between inferences made with short and long-read but particularly underscore that long-reads achieve their full potential only when paired with isoform-level analyses. Indeed, traditional event-level approaches remain severely limited, failing to capture the full scope of splicing complexity regardless of the sequencing platform used, whereas combining long-reads with isoform-centric evaluation provides the resolution necessary to uncover the true landscape of sex-biased splicing.

Using long-read technologies, we showed that sexual dimorphism in the guppy transcriptome is widespread and that males produce more isoforms in both germline and somatic tissues. While much of the elevated isoform richness in both sexes likely reflects stochastic transcriptomic noise, we find putative cases of AS driving adaptive divergence through isoform switching. These switching events are not always a strict 1:1 trade-off between male and female-biased isoforms but can also occur as complex multi-isoform dynamics from shared autosomal genes.

Ultimately, sex-biased splicing acts as a complementary process to differential gene expression as both represent routes to overcoming pleiotropic constraints and achieving sex-specific divergence.

## Methods

### Sampling and sequencing

We sampled six replicate male and female pools of two tissues, gonad and tail (muscle). All pools contained seven virgin adults for a total of 84 sampled individuals across 24 pools. RNA was extracted using Qiagen RNeasy Mini kits (Qiagen, USA) with on-column DNase following the standard protocol. RNA was quantified using an Agilent TapeStation (Agilent, USA), all RNA integrity numbers (RIN) were > 8. Samples were sent to The Centre for Applied Genomics at The Hospital for Sick Children (Toronto, Canada). Short-read sequencing was conducted using 0.8 of a lane on an Illumina NovaSeq X 25B (PE 2×150bp). Long-read sequencing (Iso-seq) was conducted using 4 SMRT cells on a PacBio Revio with 6-plex Kinnex library preparation.

### Quality control

We processed long-reads using the official PacBio IsoSeq pipeline (https://github.com/PacificBiosciences/IsoSeq) distributed through Bioconda (Grüning et al. 2018) with default settings unless otherwise noted. Briefly, we generated circular consensus sequencing (CCS) reads from subreads with *CSS* and demultiplexed using *lima*. We deconcatenated HiFi reads (Q>20) with *skera* and then demultiplexed a second time with *lima*. We trimmed poly-a tails and removed concatemers with *isoseq refine* with the --require-polya option. This resulted in the final full-length non-concatemer (FLNC) reads which we then pooled together across samples to build a common transcriptome. In total we had 195M FLNC reads with mean length 2,318bp for downstream analysis (Table S6).

We trimmed raw short-reads using *cutadapt* (Martin 2011) with --nextseq-trim command to remove poly-G reads, and *trimmomatic* to remove primers and low-quality reads (Bolger et al. 2014). After trimming we had 4.2B reads with mean length 145bp for downstream analysis.

### Building the long-read transcriptome

We clustered FLNC reads based on sequence similarity and merged to produce consensus isoforms using *isoseq cluster2*. The consensus isoforms were then mapped to the female guppy reference genome (GCF_000633615.1; Künstner et al. 2016) using *pbmm2 align*, which is the PacBio implementation of *minimap2* (Li 2018). Mapped isoforms were collapsed based on exonic structure using *isoseq collapse* with arguments --do-not-collapse-extra-5exons and –max-fuzzy-junction 0 to produce a transcriptome with unique consensus isoforms.

We used short-reads to cross-validate the long-read transcriptome with *kallisto* and *STAR* for isoform and bridge-read quantification, respectively (Dobin et al. 2013; Bray et al. 2016). We then used the output of those programs, along with the outputs from the PacBio pipeline, as input for the *SQANTI3* program for transcriptome filtering and quality control (Pardo-Palacios et al. 2024). We passed all input files to *sqanti3_qc.py* for transcriptome characterization followed by *sqanti3_filter.py* to remove isoforms with non-canonical splice sites, <10 bridge-read coverage over splice junctions, evidence of intrapriming, evidence of reverse transcriptase switching, and/or a coverage ratio of <1 at the 5’ end (Pardo-Palacios et al. 2024). We also removed all genes classified as *genic*, *genic intron*, *intergenic*, *antisense*, or *fusion* as well as all mono-exonic genes.

As in previous studies, we observed evidence of either degradation or incomplete cDNA synthesis in our long-read data despite high RIN values (Dong et al. 2023; Darolti et al. 2026). Thus, we collapsed isoforms that represented internal fragments of longer isoforms, keeping the longest one. We also set the external boundaries of all terminal junction that shared an internal boundary to match, as long-read terminal ends are known to be difficult to validate (Calvo-Roitberg et al. 2024). Although this removes our ability to identify some alternative transcription start and termination sites, it makes downstream analysis more conservative against false positives. Finally, we removed all non-coding genes as predicted by *TD2* (Mao et al. 2025) using filter psauron score <0.90, as well as all non-autosomal genes. The full long-read transcriptome construction pipeline is illustrated in Supplemental figure 8.

### Building the short-read transcriptome

In order to directly compare the results of long and short-read workflows, we built a short-read transcriptome using *STAR* and *stringtie* (Dobin et al. 2013; Pertea et al. 2015) with novel transcript discovery allowed. To match the long-read pipeline we excluded non-canonical junctions and junctions with <10 bridge-reads.

### Filtering of lowly expressed genes

To quantify isoform expression with our long-reads, we used pseudo-alignment implemented by *lr-kallisto* (Loving et al. 2025; Wissel et al. 2026) with 100 bootstraps. All further analyses were run with R in R studio unless otherwise noted (R Core Team 2025; Posit team 2026). We scaled all counts using the overdispersion coefficient calculated by catchKallisto() from the *edgeR* package, and then transformed scaled counts into TMM-normalized TPM values (Robinson et al. 2010; Baldoni et al. 2025). To measure gene-level expression, we summed isoform expression for all isoforms from a gene (Soneson et al. 2016) and retained only genes with >1 TPM in at least half of the replicates of one sex within a tissue (>=3).

We replicated this approach with the short-read transcriptome using *kallisto* and *edgeR* and filtered as above (>1 TPM in at least three samples of one sex within a tissue).

### Isoform metrics

We calculated isoform richness as the number of detected isoforms (>0 TPM) per gene. We also calculated isoform diversity, which accounts for both isoform richness and the evenness of isoform expression within a gene (isoform diversity = 2*^H^*,where *H* is the Shannon entropy of isoform expression within a gene (*H* = − ∑ *p_i_* ln *p_i_*; Schertzer et al. 2025). This measure of diversity also allows us to split isoforms into two discrete groups using an estimate of effective isoforms, which is the number of isoforms contributing substantially to gene expression given their relative expression. To do so, we determined the effective isoforms for each gene by rounding perplexity to the nearest round number *n* and then designating the *n* highest expressed isoforms as effective and all other isoforms as ineffective.

We also classified isoforms by their exon/intron structure and open-reading frame into five protein-coding potential categories to assess shifts in isoform composition between conditions. *Unique ORF*: an isoform with a complete ORF observed for the first time in the dataset*; UTR variant*: an isoform with a complete ORF that has previously been observed in the dataset and therefore differs only in UTR structure; *retained intron*: an isoform containing a retained intron and therefore unlikely to be translated due to either nonsense-mediated decay or nuclear retention (Jacob and Smith 2017); *premature STOP*: an isoform containing a premature stop codon (located >50bp upstream from the last exon-exon boundary) and therefore likely subject to nonsense-mediated decay (Kurosaki et al. 2019); *non-coding*: an isoform lacking a complete ORF, as determined by *TD2*.

### Quantifying alternative splicing (AS)

Measurement of AS between conditions can be approached in various ways. When using short-reads, AS is often quantified by measuring local splicing events which capture specific changes in exon and intron usage. Common event types include exon skipping (ES), mutually exclusive exons (MXE), alternative 5′ or 3’ splice sites (A3/A5), alternative first and last exons (AF/AL), and retained introns (RI; Figure 1A). AS is then commonly assessed using ΔPSI (delta Percent Spliced In), which measures the change in splice event inclusion between conditions, providing a quantitative estimate of differential AS.

We quantified AS at the event-level using both the long and short-read datasets with *SUPPA2* to calculate the Percent-Spliced-In (PSI) for each splicing event and tested for differential splicing between the sexes using a threshold of ΔPSI > 0.2 and p>0.05 with gene-level correction FDR < 0.05 (Trincado et al. 2018; Schertzer et al. 2025). We excluded AF/AL events given that both short and long-read technologies have difficulty assembling isoform ends reliably (Calvo-Roitberg et al. 2024).

We used the long-read dataset to quantify AS at the level of full-length isoforms. Differential isoform usage (DIU) was assessed using the diffSplice function in *edgeR*, which tests whether individual isoforms within a gene exhibit change in relative usage between conditions. Sex-biased isoform usage was evaluated by comparing male and female samples, with gene-level significance determined using the diffSplice F-test, an ANOVA-like test that assesses whether transcript-specific log-fold changes differ within a gene. Genes with FDR < 0.05 were considered to exhibit significant sex-biased splicing.

### Differential gene and isoform expression

We tested for differential gene expression (DGE) and differential isoform expression (DIE) between the sexes for both tail and gonad tissues using *edgeR* (Robinson et al. 2010; Baldoni et al. 2024). Both genes and isoforms were considered differentially expressed with an absolute log_2_ expression fold change >1 between the sexes (i.e. at least a twofold change required for significance; FDR < 0.05).

### Null model of mis-splicing

We simulated a null model of stochastic mis-splicing to estimate the isoform richness and diversity expected from transcriptional activity alone. We used an empirically estimated mis-splicing rate of 0.7% per intron (Pickrell et al. 2010) as the probability that an intron is incorrectly spliced during transcript production. For each gene, the model took as input the gene expression level and the number of introns in order to calculate the number of interactions with the spliceosome and thus opportunities for errors. We then simulated isoforms for each gene, under the assumption that the intended output is a single isoform consisting of all exons, with all other isoforms being produced as the result of mis-splicing. The output is an estimate of isoform richness and diversity per gene. Importantly, the model assumes the same mis-splicing rate across genes, sexes, and tissues and therefore provides a null expectation for isoform production driven solely by transcriptional opportunity and stochastic splicing errors.

### Identifying dynamic isoform switches between sexes

Isoform switching between the sexes is typically identified by looking for rank-order changes among the single most abundant isoform in males versus females (major isoform switching; Vitting-Seerup and Sandelin 2019) . However, this assumes that isoform switching is binary and therefore can miss coordinated changes in multiple isoforms. We opted for a broader strategy to capture greater splicing complexity. Specifically, we examined genes containing at least one male-biased and one female-biased isoform as evaluated with our DIE analysis and further required that each isoform be expressed at a biologically relevant threshold (>5 TPM). We then categorized switching events based on the number of male and female-biased isoforms produced above this threshold (1 Male:1 Female, 1 Male:Multiple Female, Multiple Male:1 Female, and Multiple Male:Multiple Female; Figure 5A).

We further classified isoform-switching genes into five categories based on the protein-coding potential of sex-biased isoforms in either sex. A*ll sex-biased ORFs*: both males and females produce distinct, sex-biased ORFs; *all redundant ORFs*: both males and females produce the same ORF but diverge in their exon structure in the UTRs; *loss of coding in one sex*: all sex-biased isoforms expressed in one sex are classified as either non-coding, premature STOP or intron retention, while the other sex retains protein-coding isoforms; *all non-coding*: all sex-biased isoforms expressed in both sexes lack protein-coding isoforms (unique ORF or UTR variant categories); *mix*: complex switching events that do not fit cleanly into the above categories.

## Data access

TBD.

## Competing interest statement

The authors declare no competing interests.

## Acknowledgements

This work was supported by an NSERC Canada Graduate Research Scholarship (Doctoral) to LMS, a Biodiversity Research Centre Biodiversity Informatics and Data Science Award to BDJ, a Birgitta Sintring Foundation Scholarship to AC-L, and a Canada 150 Research Chair and NSERC Discovery Grant to JEM. We would also like to thank Jacelyn Shu for illustrating figures 1A, 1D, 2A, 2E and 5A (jacelyndesigns.com).

## Notes

### Competing Interest Statement

The authors have declared no competing interest.

## References

Aspden JL, Wallace EWJ, Whiffin N. 2023. Not all exons are protein coding: Addressing a common misconception. Cell Genomics 3: 100296.

Ávila V, Campos JL, Charlesworth B. 2015. The effects of sex-biased gene expression and X-linkage on rates of adaptive protein sequence evolution in *Drosophila*. Biol Lett 11: 20150117.

Baldoni PL, Chen L, Li M, Chen Y, Smyth GK. 2025. Dividing out quantification uncertainty enables assessment of differential transcript usage with limma and edgeR. Nucleic Acids Res 53: gkaf1305.

Baldoni PL, Chen L, Smyth GK. 2024. Faster and more accurate assessment of differential transcript expression with Gibbs sampling and edgeR v4. NAR Genomics Bioinforma 6: lqae151.

Bénitière F, Necsulea A, Duret L. 2024. Random genetic drift sets an upper limit on mRNA splicing accuracy in metazoans. eLife 13: RP93629.

Bisazza A. 1993. Male competition, female mate choice and sexual size dimorphism in poeciliid fishes. Mar Behav Physiol 23: 257–286.

Bolger AM, Lohse M, Usadel B. 2014. Trimmomatic: a flexible trimmer for Illumina sequence data. Bioinformatics 30: 2114–2120.

Braunschweig U, Barbosa-Morais NL, Pan Q, Nachman EN, Alipanahi B, Gonatopoulos-Pournatzis T, Frey B, Irimia M, Blencowe BJ. 2014. Widespread intron retention in mammals functionally tunes transcriptomes. Genome Res 24: 1774–1786.

Bray NL, Pimentel H, Melsted P, Pachter L. 2016. Near-optimal probabilistic RNA-seq quantification. Nat Biotechnol 34: 525–527.

Burtis KC, Baker BS. 1989. Drosophila doublesex gene controls somatic sexual differentiation by producing alternatively spliced mRNAs encoding related sex-specific polypeptides. Cell 56: 997–1010.

Calvo-Roitberg E, Daniels RF, Pai AA. 2024. Challenges in identifying mRNA transcript starts and ends from long-read sequencing data. Genome Res 34: 1719–1734.

Darolti I, Labédan M, Mérel V, Schwander T. 2026. Alternative splicing shapes sexual dimorphism and erodes following the loss of sex in stick insects. http://biorxiv.org/lookup/doi/10.64898/2026.02.21.707240.

Dobin A, Davis CA, Schlesinger F, Drenkow J, Zaleski C, Jha S, Batut P, Chaisson M, Gingeras TR. 2013. STAR: ultrafast universal RNA-seq aligner. Bioinformatics 29: 15–21.

Dong X, Du MRM, Gouil Q, Tian L, Jabbari JS, Bowden R, Baldoni PL, Chen Y, Smyth GK, Amarasinghe SL, et al. 2023. Benchmarking long-read RNA-sequencing analysis tools using in silico mixtures. Nat Methods 20: 1810–1821.

Ellegren H, Parsch J. 2007. The evolution of sex-biased genes and sex-biased gene expression. Nat Rev Genet 8: 689–698.

Elliott DJ, Grellscheid SN. 2006. Alternative RNA splicing regulation in the testis. Reproduction 132: 811–819.

Fair B, Buen Abad Najar CF, Zhao J, Lozano S, Reilly A, Mossian G, Staley JP, Wang J, Li YI. 2024. Global impact of unproductive splicing on human gene expression. Nat Genet 56: 1851–1861.

Fiszbein A, Kornblihtt AR. 2017. Alternative splicing switches: Important players in cell differentiation. BioEssays 39: 1600157.

Gershoni M, Pietrokovski S. 2014. Reduced selection and accumulation of deleterious mutations in genes exclusively expressed in men. Nat Commun 5: 4438.

Gibilisco L, Zhou Q, Mahajan S, Bachtrog D. 2016. Alternative splicing within and between *Drosophila* species, sexes, tissues, and developmental stages. PLOS Genet 12: e1006464.

Gilbert W. 1978. Why genes in pieces? Nature 271: 501–501.

Grüning B, Dale R, Sjödin A, Chapman BA, Rowe J, Tomkins-Tinch CH, Valieris R, Köster J. 2018. Bioconda: sustainable and comprehensive software distribution for the life sciences. Nat Methods 15: 475–476.

Guo L, Yu H, Li Q. 2024. Sex-specific mRNA alternative splicing patterns and *Dmrt1* isoforms contribute to sex determination and differentiation of oyster. Int J Biol Macromol 283: 137747.

Harrison PW, Wright AE, Zimmer F, Dean R, Montgomery SH, Pointer MA, Mank JE. 2015. Sexual selection drives evolution and rapid turnover of male gene expression. Proc Natl Acad Sci 112: 4393–4398.

Jacob AG, Smith CWJ. 2017. Intron retention as a component of regulated gene expression programs. Hum Genet 136: 1043–1057.

Jacobs A, Elmer KR. 2021. Alternative splicing and gene expression play contrasting roles in the parallel phenotypic evolution of a salmonid fish. Mol Ecol 30: 4955–4969.

Kalsotra A, Cooper TA. 2011. Functional consequences of developmentally regulated alternative splicing. Nat Rev Genet 12: 715–729.

Karlebach G, Veiga DFT, Mays AD, Chatzipantsiou C, Barja PP, Chatzou M, Kesarwani AK, Danis D, Kararigas G, Zhang XA, et al. 2020. The impact of biological sex on alternative splicing. 490904. http://biorxiv.org/lookup/doi/10.1101/490904.

Künstner A, Hoffmann M, Fraser BA, Kottler VA, Sharma E, Weigel D, Dreyer C. 2016. The genome of the trinidadian guppy, *Poecilia reticulata*, and variation in the guanapo population. PLOS ONE 11: e0169087.

Kurosaki T, Popp MW, Maquat LE. 2019. Quality and quantity control of gene expression by nonsense-mediated mRNA decay. Nat Rev Mol Cell Biol 20: 406–420.

Li H. 2018. Minimap2: pairwise alignment for nucleotide sequences. Bioinformatics 34: 3094– 3100.

Li Y, Fang C, Fu Y, Hu A, Li C, Zou C, Li X, Zhao S, Zhang C, Li C. 2018. A survey of transcriptome complexity in *Sus scrofa* using single-molecule long-read sequencing. DNA Res 25: 421–437.

Loving RK, Sullivan DK, Reese F, Rebboah E, Sakr J, Rezaie N, Liang HY, Filimban G, Kawauchi S, Booeshaghi AS, et al. 2025. Long-read sequencing transcriptome quantification with lr-kallisto. PLOS Comput Biol 21: e1013692.

Lu Y-F, Liu Q, Liu K-Q, Wang H-Y, Li C-H, Wang Q, Shao C-W. 2022. Identification of global alternative splicing and sex-specific splicing via comparative transcriptome analysis of gonads of Chinese tongue sole (*Cynoglossus semilaevis*). Zool Res 43: 319–330.

Magurran AE, Garcia CM. 2000. Sex differences in behaviour as an indirect consequence of mating system. J Fish Biol 57: 839–857.

Mank JE. 2017. The transcriptional architecture of phenotypic dimorphism. Nat Ecol Evol 1: 0006.

Mao A, Ji HJ, Haas BJ, Salzberg SL, Sommer MJ. 2025. TD2: finding protein coding regions in transcripts. 2025.04.13.648579. http://biorxiv.org/lookup/doi/10.1101/2025.04.13.648579

Marasco LE, Kornblihtt AR. 2023. The physiology of alternative splicing. Nat Rev Mol Cell Biol 24: 242–254.

Martin M. 2011. Cutadapt removes adapter sequences from high-throughput sequencing reads. EMBnet.journal 17: 10.

Mazin PV, Khaitovich P, Cardoso-Moreira M, Kaessmann H. 2021. Alternative splicing during mammalian organ development. Nat Genet 53: 925–934.

Mi K, Guan L, Sarker B, Song S, Zhou T, Yi H, Zhang J, Xu C. 2026. Transcript diversity reflects deleterious RNA processing errors shaped by population size in metazoans. PLOS Biol 24: e3003671.

Monzó C, Liu T, Conesa A. 2025. Transcriptomics in the era of long-read sequencing. Nat Rev Genet 1–21.

Naftaly AS, Pau S, White MA. 2021. Long-read RNA sequencing reveals widespread sex-specific alternative splicing in threespine stickleback fish. Genome Res 31: 1486–1497.

Nilsen TW, Graveley BR. 2010. Expansion of the eukaryotic proteome by alternative splicing. Nature 463: 457–463.

Pardo-Palacios FJ, Wang D, Reese F, Diekhans M, Carbonell-Sala S, Williams B, Loveland JE, De María M, Adams MS, Balderrama-Gutierrez G, et al. 2024. Systematic assessment of long-read RNA-seq methods for transcript identification and quantification. Nat Methods 21: 1349–1363.

Pertea M, Pertea GM, Antonescu CM, Chang T-C, Mendell JT, Salzberg SL. 2015. StringTie enables improved reconstruction of a transcriptome from RNA-seq reads. Nat Biotechnol 33: 290–295.

Pickrell JK, Pai AA, Gilad Y, Pritchard JK. 2010. Noisy splicing drives mRNA isoform diversity in human cells. PLoS Genet 6: e1001236.

Posit team. 2026. RStudio: Integrated Development Environment for R. https://posit.co/.

R Core Team. 2025. R: A Language and Environment for Statistical Computing. https://www.R-project.org/.

Robinson MD, McCarthy DJ, Smyth GK. 2010. edgeR: a Bioconductor package for differential expression analysis of digital gene expression data. Bioinformatics 26: 139–140.

Rogers TF, Palmer DH, Wright AE. 2021. Sex-specific selection drives the evolution of alternative splicing in birds. Mol Biol Evol 38: 519–530.

Salz H, Erickson JW. 2010. Sex determination in *Drosophila*: The view from the top. Fly (Austin) 4: 60–70.

Saudemont B, Popa A, Parmley JL, Rocher V, Blugeon C, Necsulea A, Meyer E, Duret L. 2017. The fitness cost of mis-splicing is the main determinant of alternative splicing patterns. Genome Biol 18: 208.

Schertzer MD, Park SH, Su J, Reese F, Sheynkman GM, Knowles DA. 2025. Perplexity as a metric for isoform diversity in the human transcriptome. http://biorxiv.org/lookup/doi/10.1101/2025.07.02.662769.

Sharma E, Künstner A, Fraser BA, Zipprich G, Kottler VA, Henz SR, Weigel D, Dreyer C. 2014. Transcriptome assemblies for studying sex-biased gene expression in the guppy, *Poecilia reticulata*. BMC Genomics 15: 400.

Singh A, Agrawal AF. 2023. Two forms of sexual dimorphism in gene expression in *Drosophila melanogaster*: their coincidence and evolutionary genetics. Mol Biol Evol 40: msad091.

Soneson C, Love MI, Robinson MD. 2016. Differential analyses for RNA-seq: transcript-level estimates improve gene-level inferences. F1000Research 4: 1521.

Soumillon M, Necsulea A, Weier M, Brawand D, Zhang X, Gu H, Barthès P, Kokkinaki M, Nef S, Gnirke A, et al. 2013. Cellular source and mechanisms of high transcriptome complexity in the mammalian testis. Cell Rep 3: 2179–2190.

Steijger T, Abril JF, Engström PG, Kokocinski F, Hubbard TJ, Guigó R, Harrow J, Bertone P. 2013. Assessment of transcript reconstruction methods for RNA-seq. Nat Methods 10: 1177–1184.

Su Y, Yu Z, Jin S, Ai Z, Yuan R, Chen X, Xue Z, Guo Y, Chen D, Liang H, et al. 2024. Comprehensive assessment of mRNA isoform detection methods for long-read sequencing data. Nat Commun 15: 3972.

Telonis-Scott M, Kopp A, Wayne ML, Nuzhdin SV, McIntyre LM. 2009. Sex-specific splicing in *Drosophila*: Widespread occurrence, tissue specificity and evolutionary conservation. Genetics 181: 421–434.

Tosto NM, Beasley ER, Wong BBM, Mank JE, Flanagan SP. 2023. The roles of sexual selection and sexual conflict in shaping patterns of genome and transcriptome variation. Nat Ecol Evol 7: 981–993.

Trincado JL, Entizne JC, Hysenaj G, Singh B, Skalic M, Elliott DJ, Eyras E. 2018. SUPPA2: Fast, accurate, and uncertainty-aware differential splicing analysis across multiple conditions. Genome Biol 19: 40.

van der Bijl W, Shu JJ, Goberdhan VS, Sherin LM, Jia C, Cortazar-Chinarro M, Corral-Lopez A, Mank JE. 2025. Deep learning reveals the complex genetic architecture of male guppy colouration. Nat Ecol Evol 9: 1614–1625.

Verta J-P, Jacobs A. 2022. The role of alternative splicing in adaptation and evolution. Trends Ecol Evol 37: 299–308.

Vitting-Seerup K, Sandelin A. 2019. IsoformSwitchAnalyzeR: Analysis of changes in genome-wide patterns of alternative splicing and its functional consequences. Bioinformatics 35: 4469–4471.

Wan Y, Larson DR. 2018. Splicing heterogeneity: separating signal from noise. Genome Biol 19: 86.

Weatheritt RJ, Sterne-Weiler T, Blencowe BJ. 2016. The ribosome-engaged landscape of alternative splicing. Nat Struct Mol Biol 23: 1117–1123.

Wissel D, Mehlferber MM, Nguyen KM, Pavelko V, Tseng E, Robinson MD, Sheynkman GM. 2026. A systematic benchmark of high-accuracy PacBio long-read RNA sequencing for transcript-level quantification. Genome Biol 27: 110.

Wright CJ, Smith CWJ, Jiggins CD. 2022. Alternative splicing as a source of phenotypic diversity. Nat Rev Genet 23: 697–710.

