## Supplemental figures 1-8 for "Elevated isoform richness in males largely reflects transcriptional noise rather than proteomic complexity"

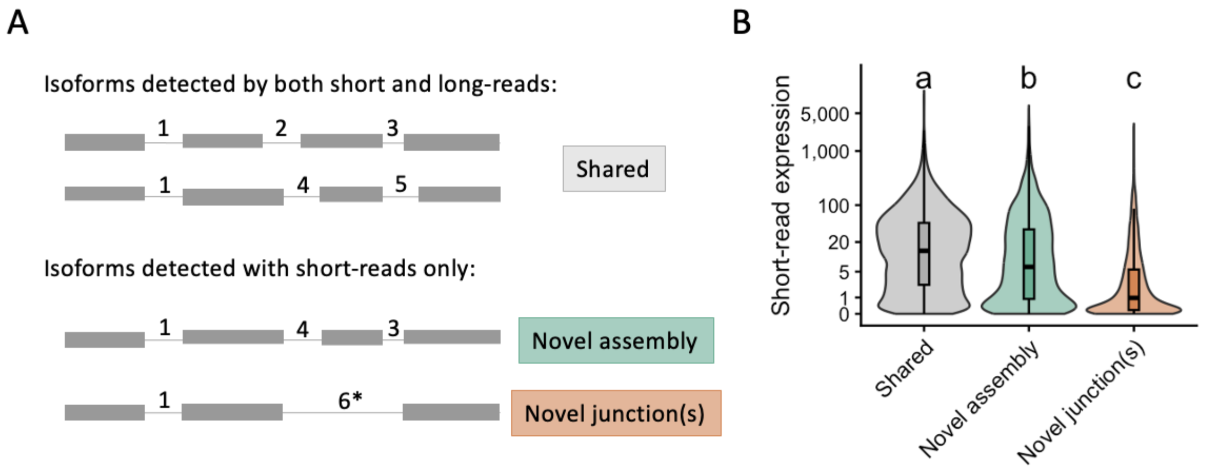


***Supplemental figure 1:*** (A) Isoforms that are unique to the short-read dataset may either be novel assemblies of known splice junctions (green) or assembles including novel junctions not recovered in the long-read dataset (orange). (B) The isoforms with novel junctions are found to have lower expression ($log_2$(TPM+1), supporting the idea that they represent isoforms either not detected or filtered out of the long-read transcriptome due to insufficient read depth. Letters above boxplots indicate statistically significant differences among groups based on Kruskal-Wallis tests followed by Dunn's post hoc pairwise comparisons with Benjamini-Hochberg correction for multiple testing (FDR < 0.05).

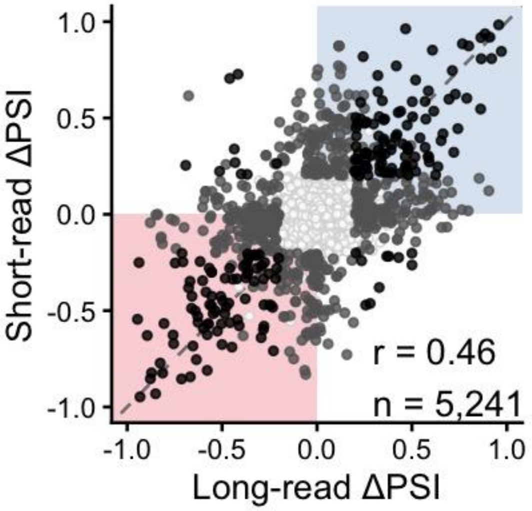


***Supplemental figure 2:*** The difference in percent-spliced-in (ΔPSI) between males and females as measured by the long and short-read datasets, coloured by significance. Plotted here are only those events found in both datasets. Significance requires a p-value <0.05 and ΔPSI > 0.2 (20% difference in expression between conditions). Blue quadrant indicates male-biased in both datasets, red quadrant indicates female-biased in both datasets. Pearson correlation, *r* = 0.46, p < 0.001. Correlation (*r*) and number of observations (n) are annotated on lower right-hand quadrant.

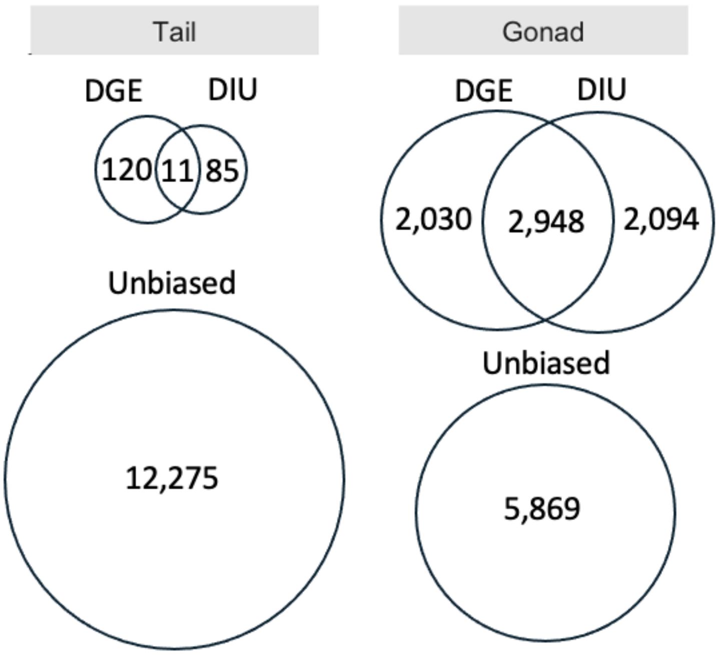


***Supplemental figure 3:*** Venn diagrams showing the number of genes found to exhibit differential gene expression (DGE) and differential isoform usage (DIU; sex-biased splicing). In tail, most genes were neither significantly sex-biased in expression or alternatively spliced between the sexes. We observed much more transcriptional complexity in the gonad as 22.8% of genes in the gonad exhibited both sex-biased gene expression and alternative splicing, and a further 16.2% were unbiased in their gene expression but alternatively spliced between the sexes.

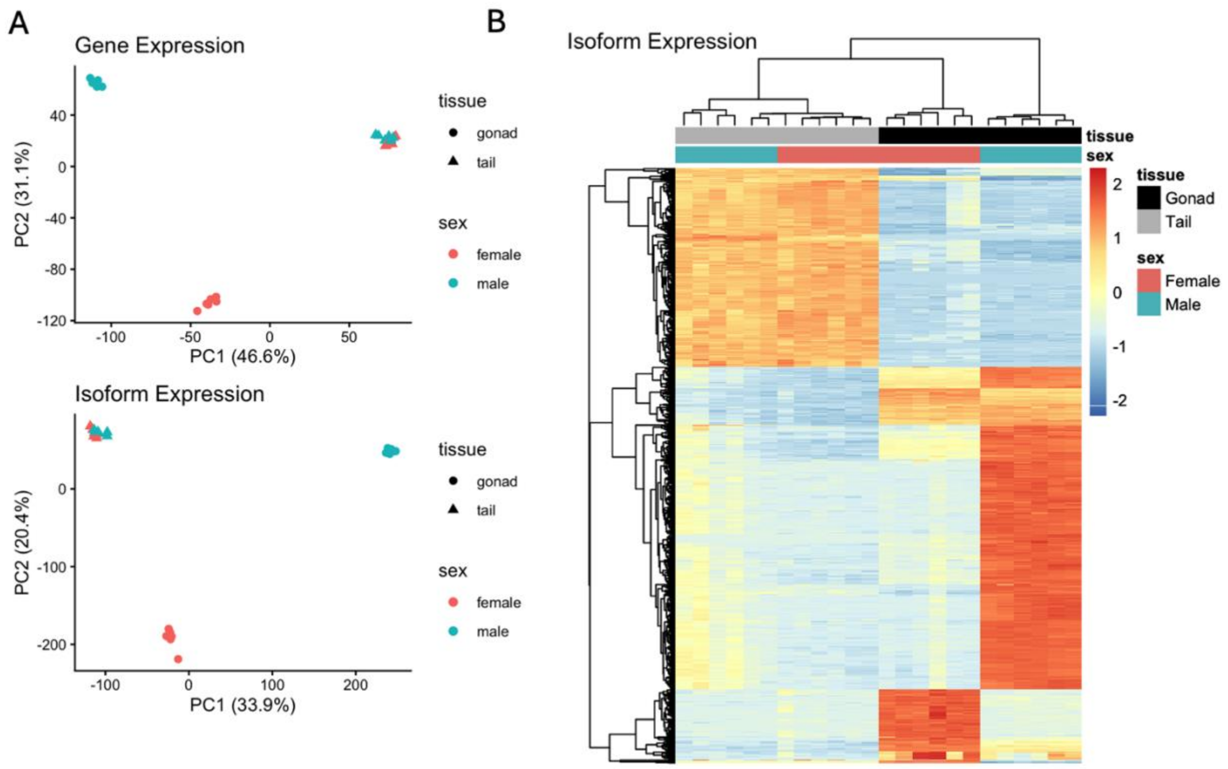


***Supplemental figure 4:*** (A) PCA of gene expression for all 24 samples. (B) PCA of isoform expression. (D) Heatmap of the expression of 1000 most variable isoforms with hierarchical clustering of samples (x-axis) and genes (y-axis). Expression is in units log2(TPM + 1) for all panels.

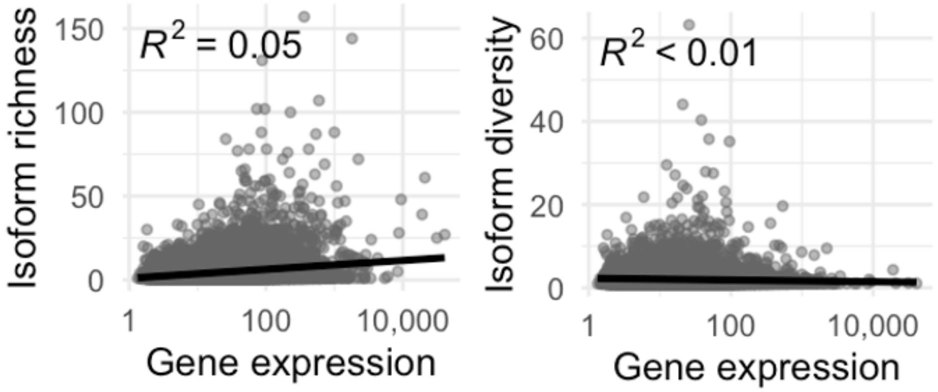


***Supplemental figure 5:*** (Left) Isoform richness has a positive relationship with gene expression level (log2(TPM+1)). (Right) Isoform diversity has no relationship with gene expression level. X-axis is shown on a log scale. Both plots show metrics calculated after aggregating data from all samples across sex and tissues.

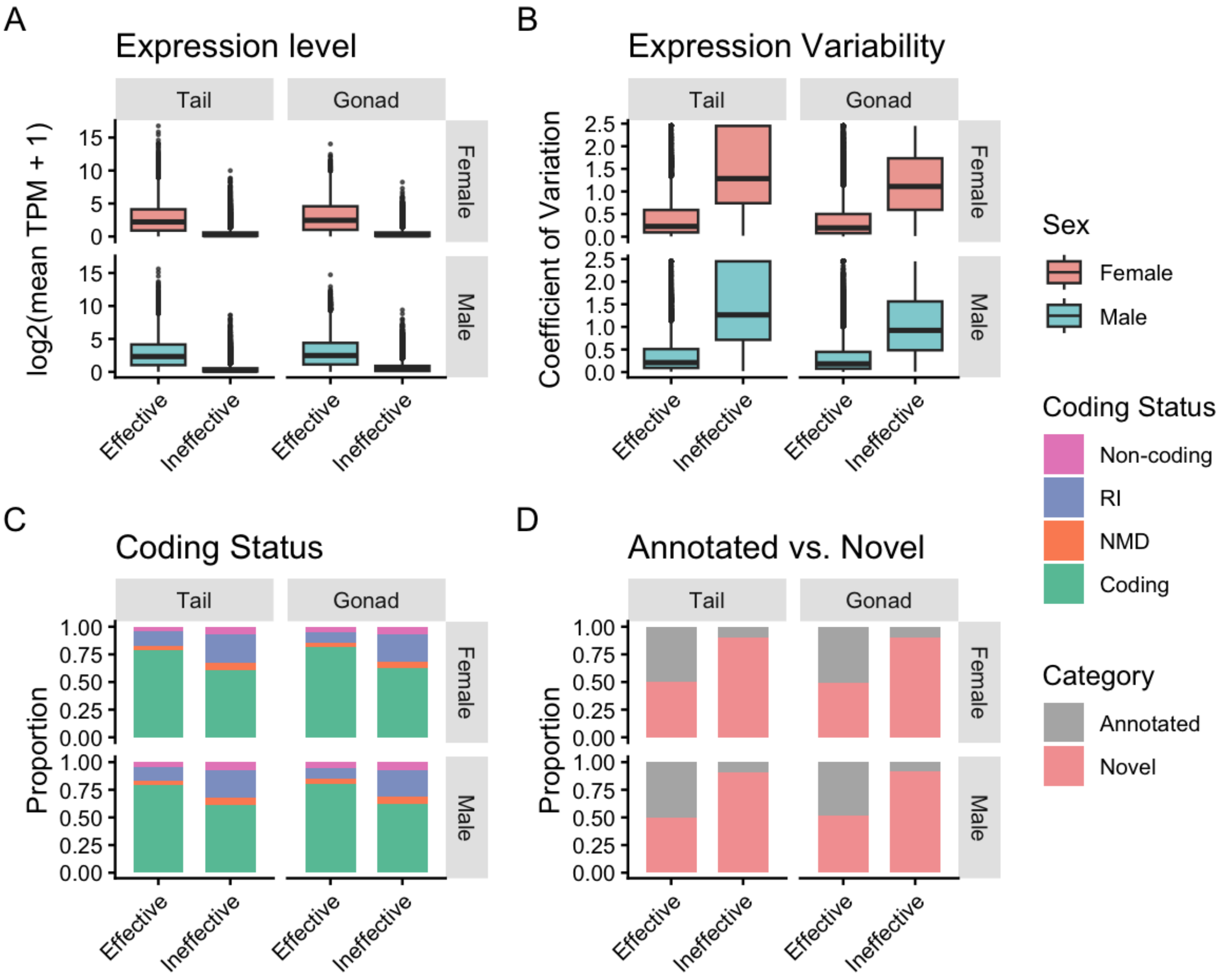


***Supplemental figure 6:*** (A) Isoform expression level measured as log2(mean TPM + 1). Effective isoforms generally show higher expression than ineffective isoforms. (B) Expression variability measured as the coefficient of variation across samples. Ineffective isoforms exhibit greater expression variability than effective isoforms in both tissues and sexes. (C) Coding status of effective and ineffective isoforms, showing the proportions classified as coding, nonsense-mediated decay (NMD), retained intron (RI), and non-coding by SQANTI3. Effective isoforms are more often coding, whereas ineffective isoforms contain higher proportions of non-coding, NMD and RI isoforms. (D) Proportion of annotated and novel isoforms as classified by SQANTI3. Effective isoforms contain a larger fraction of annotated transcripts, while ineffective isoforms tend to be unannotated (novel). Results are shown separately for tail and gonad tissues and for female and male samples.

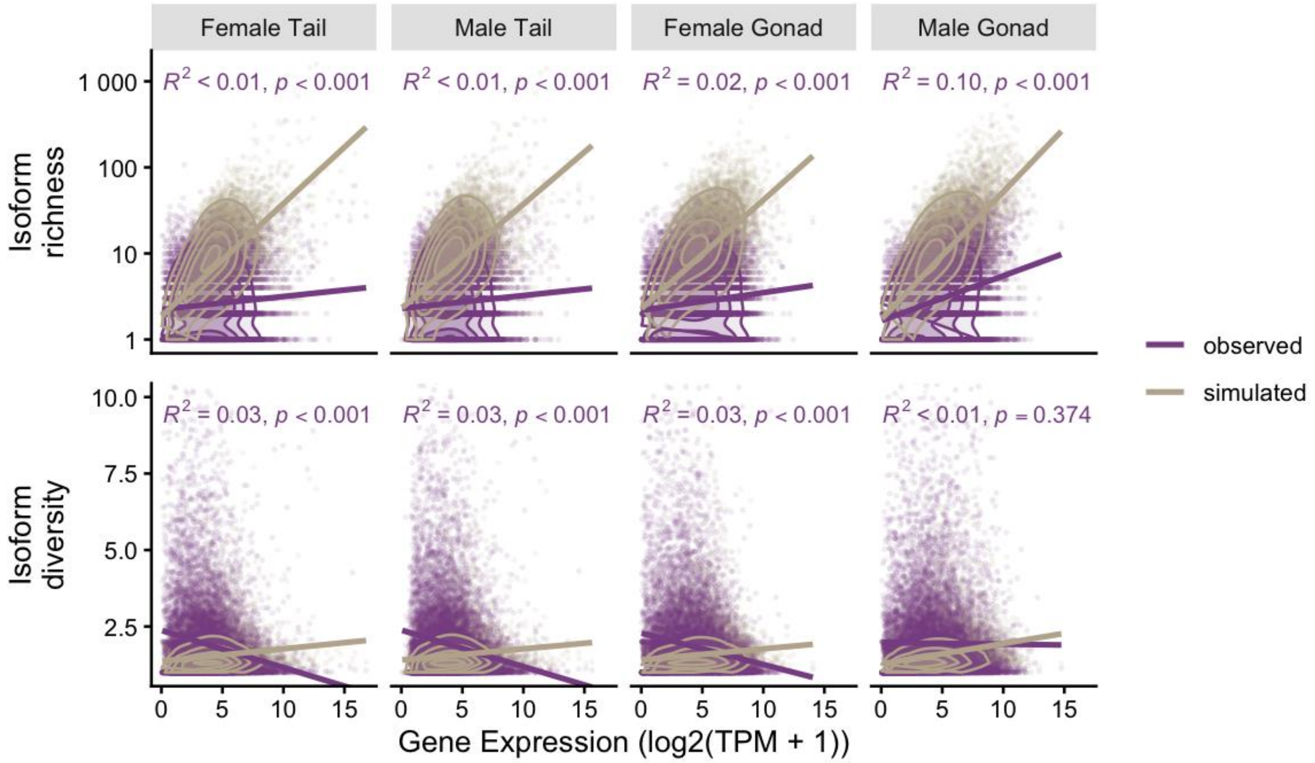


***Supplemental figure 7:*** Comparison of observed relationships of gene expression and isoform richness and diversity with a null model of mis-splicing. The simulation assumes a mis-splicing rate of 0.7% per intron (Pickrell et al. 2010) and takes as input the gene expression level of a gene and the number of introns which together represent the number of interactions with the spliceosome. The output is isoform richness and diversity per gene. Top row: Isoform richness, values shown on a log scale. Bottom row: Isoform diversity.

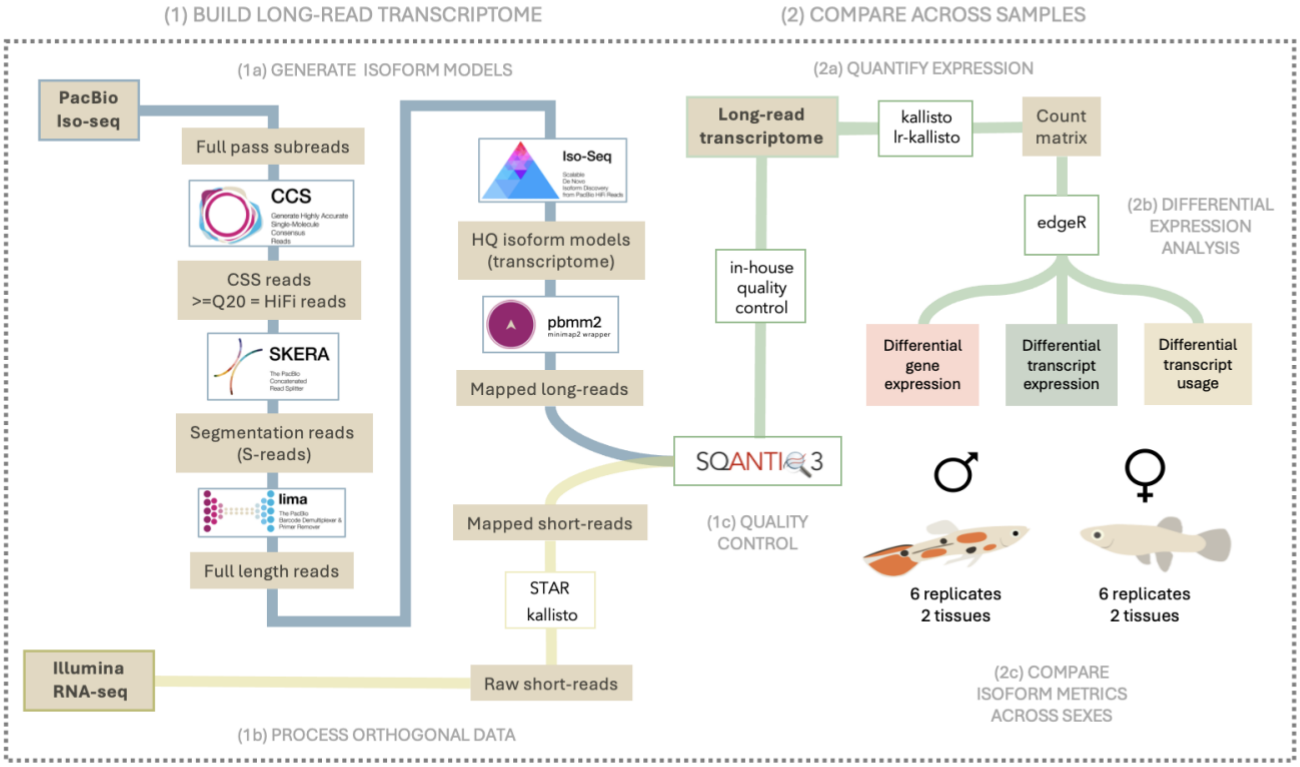


***Supplemental figure 8:*** Workflow to construct the long-read transcriptome using short-read orthogonal data, filter the transcriptome to only highly supported isoform models and finally quantify expression and other isoform metrics to conduct comparative analyses between the sexes.
